# Defining the Regulatory Logic of Breast Cancer Using Single-Cell Epigenetic and Transcriptome Profiling

**DOI:** 10.1101/2024.06.13.598858

**Authors:** Matthew J. Regner, Susana Garcia-Recio, Aatish Thennavan, Kamila Wisniewska, Raul Mendez-Giraldez, Brooke Felsheim, Philip M. Spanheimer, Joel S. Parker, Charles M. Perou, Hector L. Franco

## Abstract

Annotation of the *cis*-regulatory elements that drive transcriptional dysregulation in cancer cells is critical to improving our understanding of tumor biology. Herein, we present a compendium of matched chromatin accessibility (scATAC-seq) and transcriptome (scRNA-seq) profiles at single-cell resolution from human breast tumors and healthy mammary tissues processed immediately following surgical resection. We identify the most likely cell-of-origin for luminal breast tumors and basal breast tumors and then introduce a novel methodology that implements linear mixed-effects models to systematically quantify associations between regions of chromatin accessibility (i.e. regulatory elements) and gene expression in malignant cells versus normal mammary epithelial cells. These data unveil regulatory elements with that switch from silencers of gene expression in normal cells to enhancers of gene expression in cancer cells, leading to the upregulation of clinically relevant oncogenes. To translate the utility of this dataset into tractable models, we generated matched scATAC-seq and scRNA-seq profiles for breast cancer cell lines, revealing, for each subtype, a conserved oncogenic gene expression program between *in vitro* and *in vivo* cells. Together, this work highlights the importance of non-coding regulatory mechanisms that underlie oncogenic processes and the ability of single-cell multi-omics to define the regulatory logic of BC cells at single-cell resolution.

## INTRODUCTION

Breast cancer (BC) is the most commonly diagnosed cancer among women and accounts for 15% of all female cancer-related deaths in the United States(1). Treatment strategies and patient prognosis vary by clinical subtype, defined by hormone receptor expression of estrogen receptor (ER), progesterone receptor (PR), and overexpression and/or amplification of human epidermal growth factor receptor 2 (HER2). BC can also be stratified into five intrinsic molecular subtypes with distinct clinical outcomes: Luminal A, Luminal B, HER2-enriched, Basal-like, and Normal-like(2-5). Together, these form three broad subtypes of BC: luminal (ER^+^/PR^+/-^), HER2+ (HER2^+^, ER^+/−^, PR^+/−^), and triple-negative (ER^-^, PR^-^, HER2^-^) BC(6-8). Several studies have characterized the transcriptional landscapes of these BC subtypes(9-12). While these studies have been transformative, there has been an increased focus on the non-coding regions of the genome, in addition to transcriptomics, for deeper multi-omic insights into BC heterogeneity and its pathogenesis(13-16).

Non-coding regions contain vast amounts of regulatory information that contribute profoundly to tumor biology. Moreover, it has become increasingly evident that regulatory elements (i.e. *cis*-acting enhancers) are re-wired in cancer cells to promote growth, survival, and other aggressive phenotypes associated with poor clinical outcome(17-25). Several studies have used epigenomics, in parallel with transcriptomics, to characterize the molecular and clinical heterogeneity of BC revealing extensive variation in enhancer activity across BC subtypes(13-16). However, most studies to date have done so using bulk genomic sequencing of material collected from heterogenous mixtures of different cell types, obscuring the cancer cell-specific activity of oncogenic enhancers. Therefore, the exact mechanisms of gene regulation in the context of BC cells remain elusive.

Breast tumors are complex cellular microenvironments in which various types of malignant and non-malignant cells contribute to a range of clinically relevant biological phenomena, from cancer progression to treatment response(26-28). It is widely accepted that BC arises from mammary epithelial cells(29, 30). The normal mammary epithelium is mainly comprised of mature luminal, luminal progenitor, and basal epithelial cells, all of which have been studied as possible “cell-of-origin” precursors for different BC molecular subtypes(31-35). Several studies have proposed luminal progenitor and mature luminal cells as the likely cell-of-origin precursors for Basal-like and Luminal BC, respectively (31-35). However, changes in the gene regulatory landscape between normal mammary epithelial and subtype-specific BC cells, are not as well studied, especially at single-cell resolution.

Single-cell genomics has revolutionized our ability to explore the cellular heterogeneity of breast tumors, yet most studies have profiled transcriptomes via single-cell RNA-seq (scRNA-seq)(28, 36-42). The single-cell assay for transposase-accessible chromatin by sequencing (scATAC-seq) performs high-throughput profiling of chromatin accessibility, revealing complex facets of gene regulation, including the activity of enhancers at single-cell resolution(43-45). Together, scRNA-seq and scATAC-seq enable the linking of regulatory elements to putative target genes, offering key mechanistic insights into the molecular underpinnings of BC by interrogating the regulatory logic of BC cells(23, 35, 46-52). We posit that enhancers with increased activity in BC cells, relative to normal mammary epithelial cells, regulate the expression of genes associated with oncogenic processes.

To investigate how regulatory landscapes may become altered in human breast cancers relative to the normal mammary epithelium, we generated matched scRNA-seq and scATAC-seq profiles for twelve primary breast tumor specimens, four normal breast tissue specimens, and four BC cell lines. This compendium of matched scRNA-seq and scATAC-seq data, encompassing over 200,000 single cells from breast tumor tissues, true normal mammary tissues, and BC cell lines, will serve as an important resource to the single-cell genomics and BC research communities. Moreover, we introduce a novel methodology that implements linear mixed-effects models (LMMs) to quantify associations between regions of chromatin accessibility (i.e. regulatory elements) and gene expression while accounting for important biological and technical variables(53, 54).

The LMM-based approach enabled us to perform differential association analyses between subtype-specific BC cells and their nearest normal mammary epithelial cells from healthy controls (e.g., comparing Basal-like BC to luminal progenitor and Luminal BC to mature luminal). We also apply the LMM-based method to BC cell lines and compare the regulatory landscapes between BC cells *in vivo* and *in vitro*, stratified by molecular subtype. Through these analyses, we identify context-specific mechanisms of gene regulation in BC cells and unveil clinically relevant non-coding mechanisms for BC pathogenesis at single-cell resolution.

## RESULTS

### Matched scRNA-seq and scATAC-seq of human breast tumors and normal mammary epithelial tissues

Twelve primary breast tumor specimens and four normal mammary tissue specimens were collected from eleven treatment-naïve BC patients undergoing surgery with curative intent and four healthy control patients undergoing a reduction mammoplasty procedure, respectively (Table 1; Table S1; Figure 1). Two specimens were collected from BC Patient 14 (Table 1; Table S1). Immediately following surgical resection, each specimen was dissociated into a live cell suspension and prepared for droplet-based scRNA-seq and scATAC-seq on the 10X Genomics Chromium platform (Figure 1A; Methods). After quality control for each patient dataset, we obtained 111,879 cells and 91,048 cells profiled by scRNA-seq and scATAC-seq, respectively (Table 1; Table S2A; Table S3A; Figure S1).

**Table 1.** Abbreviated clinical data and single-cell metadata for each sample.

| Sample | Type | ER IHC | PR IHC | Her2 IHC | scRNA-seq cells | scATAC-seq cells |
| --- | --- | --- | --- | --- | --- | --- |
| Patient 1 | Normal Breast Tissue | Not Performed | Not Performed | Not Performed | 8,682 | 8,166 |
| Patient 2 | Normal Breast Tissue | Not Performed | Not Performed | Not Performed | 6,971 | 9,337 |
| Patient 3 | Normal Breast Tissue | Not Performed | Not Performed | Not Performed | 6,368 | 9,549 |
| Patient 4 | Normal Breast Tissue | Not Performed | Not Performed | Not Performed | 10,222 | 6,860 |
| Patient 5 | Primary Breast Tumor | - | + | - | 6,583 | 5,647 |
| Patient 6 | Primary Breast Tumor | - | - | - | 3,965 | 1,429 |
| Patient 7 | Primary Breast Tumor | + | - | - | 7,029 | 4,093 |
| Patient 8 | Primary Breast Tumor | + | + | Equivocal | 10,085 | 6,490 |
| Patient 9 | Primary Breast Tumor | + | + | Equivocal | 6,907 | 2,025 |
| Patient 10 | Primary Breast Tumor | + | + | Equivocal | 5,307 | 5,016 |
| Patient 11 | Primary Breast Tumor | + | + | - | 2,997 | 2,743 |
| Patient 12 | Primary Breast Tumor | + | + | Equivocal | 8,490 | 6,339 |
| Patient 13 | Primary Breast Tumor | + | + | - | 6,676 | 3,383 |
| Patient 14-1 | Primary Breast Tumor | + | + | - | 9,175 | 8,010 |
| Patient 14-2 | Primary Breast Tumor | + | + | - | 5,655 | 4,768 |
| Patient 15 | Primary Breast Tumor | + | + | Equivocal | 6,767 | 7,193 |
| MCF7 | Breast Cancer Cell Line | Not Performed | Not Performed | Not Performed | 7,331 | 4,367 |
| T47D | Breast Cancer Cell Line | Not Performed | Not Performed | Not Performed | 7,796 | 6,709 |
| HCC1143 | Breast Cancer Cell Line | Not Performed | Not Performed | Not Performed | 6,827 | 1,313 |
| SUM149PT | Breast Cancer Cell Line | Not Performed | Not Performed | Not Performed | 8,697 | 3,949 |

**Figure 1.**
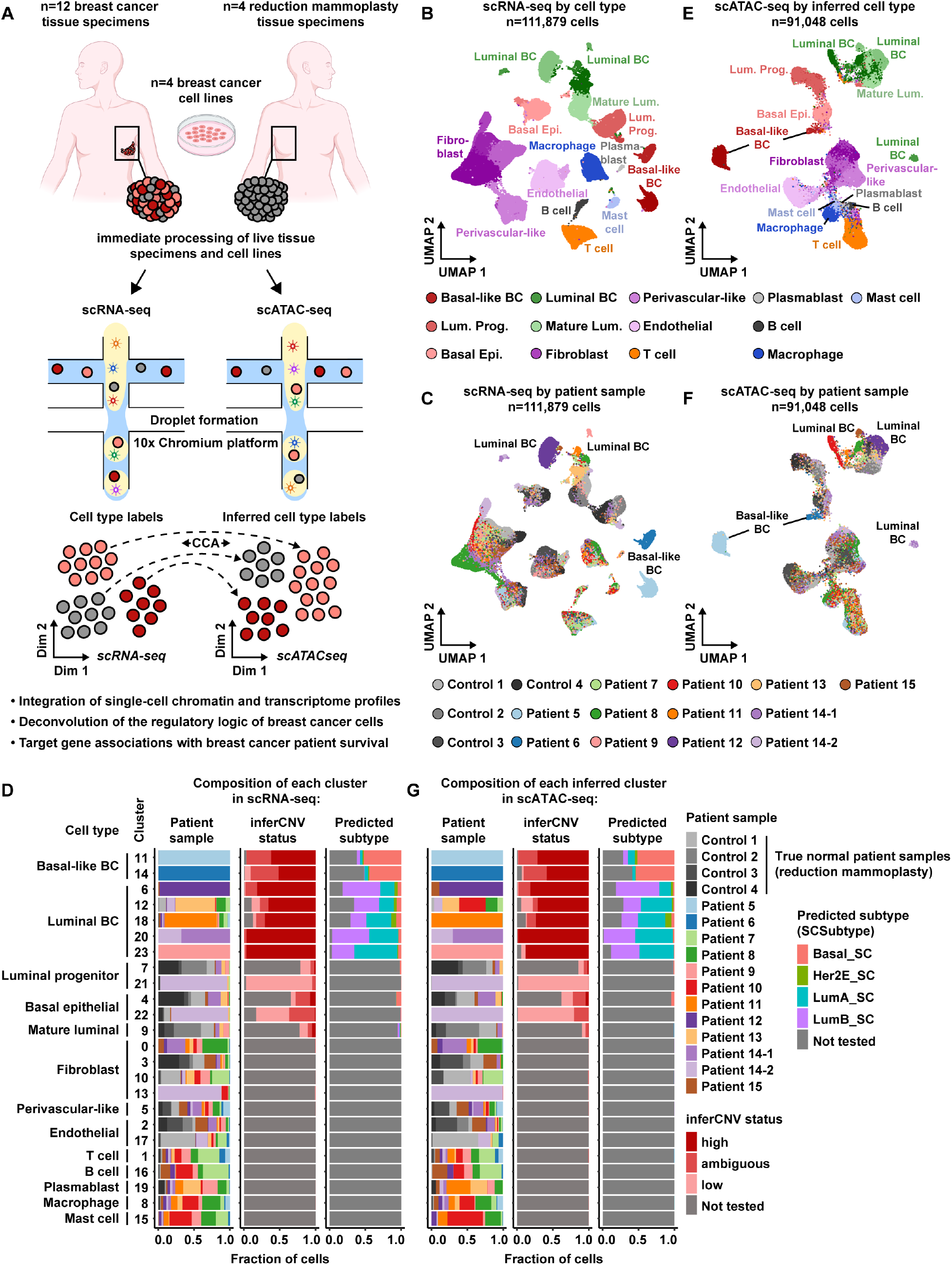
Overview of matched scRNA-seq and scATAC-seq workflow for BC tissue specimens, reduction mammoplasty tissue specimens, and BC cell lines. **A)**. Schematic diagram of the procurement, processing, and downstream analysis of patient samples and cell lines. The female breast and cell line illustrations were created with BioRender.com. **B)**. UMAP plot of 111,879 scRNA-seq cells color-coded by cell type across 16 patient samples. Color shades denote clusters within each cell type**. C)**. UMAP plot of scRNA-seq cells, as shown in panel **B**, but color-coded by the patient sample of origin. **D)**. Proportion bar charts showing the composition of each cluster in scRNA-seq, in terms of patient sample (*left*), inferCNV status (*middle*), and predicted subtype (*right*). Color-coded legends are shown to the right. **E)**. UMAP plot of 91,048 scATAC-seq cells color-coded by inferred cell type across 16 patient samples. Color shades denote clusters within each inferred cell type**. F)**. UMAP plot of scATAC-seq cells, as shown in panel **E**, but color-coded by the patient sample of origin. **G)**. Proportion bar charts, as in panel **D**, but for the composition of each inferred cluster in scATAC-seq.

To analyze the scRNA-seq cells from all 16 patient samples, we used the top 2,000 most variably expressed genes across all 111,879 cells to perform principal component analysis (PCA). Cells were clustered using graph-based clustering, with the top 30 principal components (PCs), and visualized in a uniform manifold approximation and projection (UMAP) plot. This showed clusters that were annotated to known cell types (Figure 1B; Figure S2A-B; Table S2A; Methods) and that batch effects were not a major source of variation (Figure 1C; Figure 1D)(28, 46, 47).

To identify malignant BC cells within each patient tumor, we used inferCNV to estimate copy-number variant (CNV) profiles at single-cell resolution as described previously(28, 55-57). Briefly, this procedure involved classifying epithelial cells from each patient tumor into one of three groups (high, ambiguous, or low) based on the inferred level of CNV in each cell(28). Cells classified as inferCNV high were deemed putative cancer cells and were carried forward to molecular subtype prediction. We predicted the molecular subtype (Basal-like, Her2-enriched, Luminal A, or Luminal B) of each cancer cell within each patient tumor using a previously published method called SCSubtype(28). These analyses revealed the majority compositions of clusters 11 and 14 as Basal-like BC cells from Patients 5 and 6, respectively (Figure 1D; Figure S2B; Table S2A). We observed that the majority compositions of clusters 6, 12, 18, 20, and 23, from Patients 12, 13, 11, 14, and 9, respectively, contained mixtures of predicted Luminal A and B cells, suggesting these cells can be referred to hereafter as Luminal breast cancer (Figure 1D; Figure S2B; Table S2A). We finally observed that the normal mammary epithelial cell type clusters of luminal progenitor, basal epithelial, and mature luminal cells were well-represented by the healthy control samples, indicating a sufficient number of cells for a robust control group comparison to BC cells (Figure 1D; Figure S2A-B; Table S2A).

To analyze the scATAC-seq cells from all 16 patient samples, we first quantified Tn5 insertion counts in a matrix of contiguous genomic tiles before applying iterative latent semantic indexing with the top 10,000 most variably accessible genomic tiles to reduce the dimensionality of the dataset (44, 45, 48). Using Seurat’s cross-modality integration approach, we then assigned cell type cluster labels to scATAC-seq cells based on their matching scRNA-seq data (Figure 1E; Figure 1G; Figure S2C-E; Table S3A; Methods)(46-48). The cross-modality integration approach also enabled us to assign inferCNV status and predicted subtype to each scATAC-seq cell based on the annotations of its nearest neighboring cell in scRNA-seq (Figure 1G; Table S3A;)(46-48). This showed that the scATAC-seq cells clustered mainly by cell type, not by patient, consistent with the matching scRNA-seq data (Figure 1F; Figure 1G). In summary, we observed 13 general cell types across the patient dataset, with 24 clusters identified in both scRNA-seq and scATAC-seq. Consistent with previous reports, we observed that the non-malignant cell type clusters were well-represented across patients, while the BC clusters remained highly patient-specific (Figure 1D; Figure S2B)(23, 28, 58-60).

To further inform our comparisons of subtype-specific BC cells from primary tumors to normal mammary epithelial cells from healthy controls, we performed an unsupervised clustering analysis of pseudo-bulk transcriptomes (Figure S3). To create BC cell-specific pseudo-bulk transcriptomes, we summed counts across BC cells of the majority subtype within each BC patient sample (Figure 1D; Figure S2B). The same was performed for each healthy control’s normal mammary epithelial cell types (Figure 1D; Figure S2B). Unsupervised hierarchical clustering of these pseudo-bulk transcriptional profiles revealed the Luminal BC profiles from BC patients clustered with mature luminal profiles from healthy controls, while Basal-like profiles from BC patients clustered with luminal progenitor and basal epithelial profiles from healthy controls (Figure S3A). To further investigate this, we performed PCA of the same pseudo-bulk transcriptional profiles, which revealed clear separation of the Basal-like BC, luminal progenitor, and basal epithelial profiles from the Luminal BC and mature luminal profiles (Figure S3B). These observations were consistent with previous reports supporting luminal progenitor and mature luminal cells as the cell-of-origin precursors to Basal-like and Luminal BC, respectively(31, 32, 35). To this end, we transitioned into subtype-specific analyses of BC (Basal-like and Luminal BC) compared to their nearest normal mammary epithelial cell types.

### Identification of enhancers with cancer-specific regulatory activity in Basal-like BC cells

Basal-like BC has been shown to strongly overlap with the triple-negative clinical subtype, and portends a poor prognosis, in part, due to a lack of targeted therapies(61-66). To analyze the Basal-like subtype cells, we merged Basal-like BC cells from Patients 5 and 6 with luminal progenitor as well as basal epithelial cells from the healthy mammary reduction patients, according to the results of the unsupervised pseudo-bulk clustering analysis (Figure S3). This subset resulted in 13,993 cells and 14,038 cells profiled by scRNA-seq and scATAC-seq, respectively (Figure 2A; Table S2B; Table S3B). After re-clustering the scRNA-seq cells and transferring the resulting labels as well as gene expression profiles to the scATAC-seq cells, we observed that cells mainly clustered by cell type and not by patient, except for two patient-specific clusters of Basal-like BC from Patients 5 and 6 (Figure 2B; Table S2B; Table S3B).

**Figure 2.**
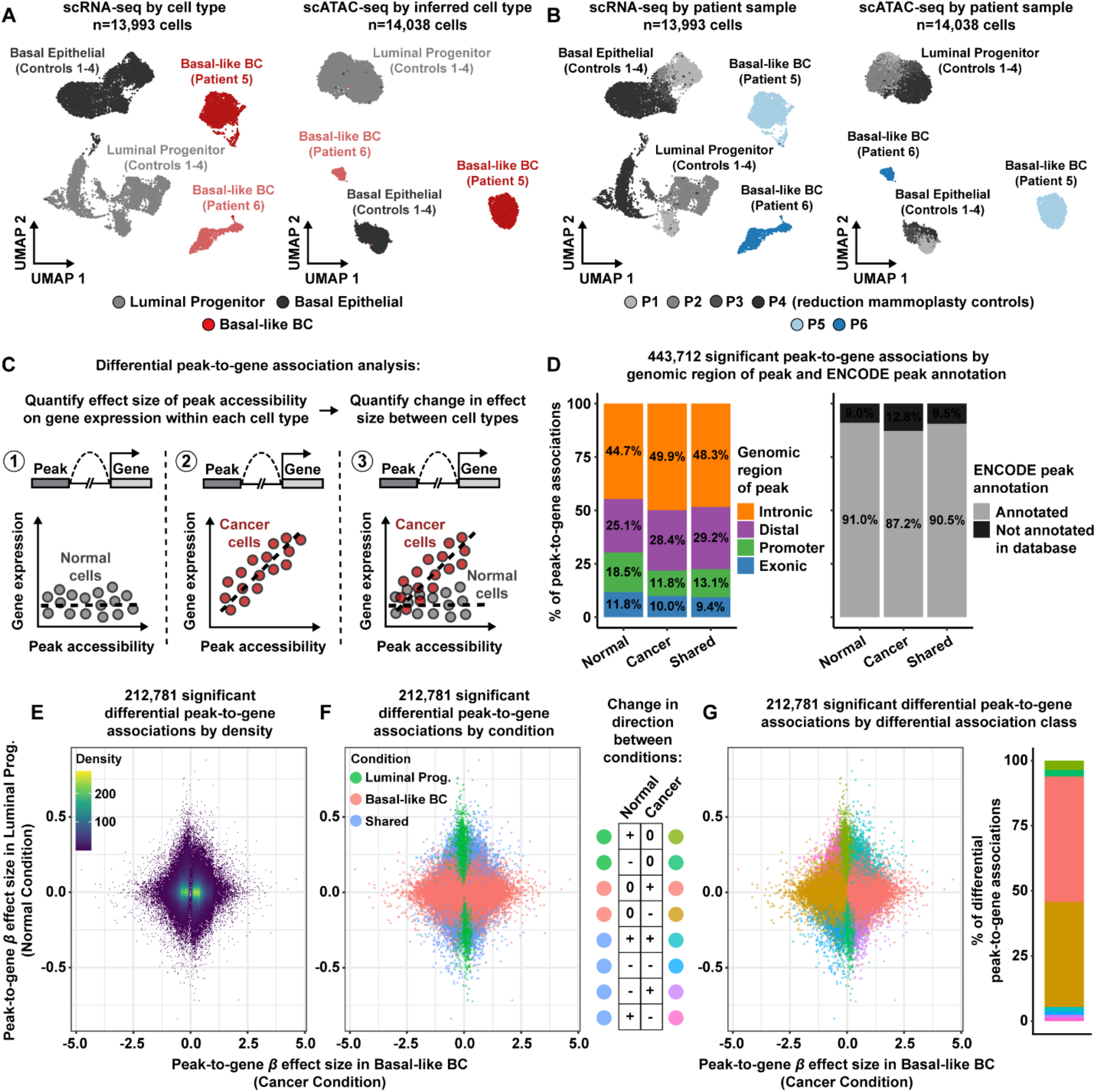
Quantifying the altered regulatory landscape in Basal-like BC cells relative to normal luminal progenitor cells. **A).** UMAP plot of 13,993 scRNA-seq cells color-coded by cell type across 6 patient samples (*left*). UMAP plot of 14,038 scATAC-seq cells color-coded by inferred cell type across 6 patient samples (*right*). Color shades denote clusters within each cell type**. B).** UMAP plots of scRNA-seq cells (*left*) and scATAC-seq cells (*right*), as shown in panel **A**, but color-coded by the patient sample of origin. **C).** Schematic diagram showing the differential peak-to-gene analysis framework in three steps. Quantifying the effect size of chromatin accessibility on the inferred level of gene expression for a given peak-gene pair, in normal (*left*) and in cancer cells (*middle*). Quantifying the change in effect size of chromatin accessibility on the inferred level of gene expression for the same peak-gene pair between normal and cancer cells (*right*). **D).** Proportion bar charts showing the genomic distribution (*left*) and ENCODE annotation status (*right*) for 84,975 normal-specific peak-to-gene associations, 337,053 cancer-specific associations, and 21,684 shared associations. **E).** Density scatter plot showing the effect sizes of significant differential peak-to-gene associations in the cancer condition, comprised of Basal-like BC cells, and in the normal condition, comprised of luminal progenitor cells. Each dot represents a peak-gene pair with a significant change in effect size between conditions. **F).** Same scatter plot as in panel **E**, but colored by condition specificity (*left*). Infographic describing the possible directional changes in effect size between conditions (*right*). **G).** Same scatter plot as in panel **F**, but colored by changes in direction of effect size between conditions (*left*). Proportion bar chart showing the distribution of differential association classes (*right*).

To interrogate the regulatory logic of Basal-like BC cells in comparison to their nearest normal mammary epithelial cells, we first carried out peak calling in scATAC-seq cells to identify putative regulatory elements located in regions of accessible chromatin(13, 48, 67, 68). We developed a robust LMM-based strategy to link putative regulatory elements to target genes. This enabled us to quantify the association between peak accessibility and gene expression after accounting for patient. To investigate the regulatory landscapes in Basal-like BC cells compared to normal mammary epithelial cells, we performed a two-phased differential peak-to-gene association analysis to link putative regulatory elements to target genes in a context-specific manner (Figure 2C; Methods).

Consistent with previous reports, we focused our analysis on the comparison of Basal-like BC cells, referred to hereafter as the “cancer” condition, to luminal progenitor cells, referred to hereafter as “normal” (31, 32, 35). Briefly, we identified groups of similar scATAC-seq cells within each patient via k-nearest neighbors and aggregated their sparse profiles into metacell observations for the peak-to-gene association analysis (48). Using the patient-specific metacells, we first quantified, within each condition, the regulatory effect size of peak accessibility on gene expression for every peak within 500 kb of each gene, after accounting for variation between patients (Figure 2C)(53, 54). More specifically, for every peak-gene pair tested in each condition, we modeled gene expression as a function of peak accessibility as a fixed effect and patient as a random effect in an LMM. This revealed a total of 443,712 significant peak-to-gene associations (FDR-adjusted p-value < 1e-04), with 84,975 normal-specific associations, 337,053 cancer-specific associations, and 21,684 shared associations (Figure 2D; Table S4). We observed that the majority of these peak-to-gene associations involved peaks located in introns and distal intergenic regions, highlighting the importance of non-coding regulatory information (Figure 2D; Table S4)(23). Moreover, the majority of these peak-to-gene associations also involved peaks annotated by the Encyclopedia of DNA Elements Consortium (ENCODE) database, suggesting they are bona fide regulatory elements that provide support for our computational approach (Figure 2D; Table S4)(69, 70).

To identify putative regulatory elements with significant differential effects on gene expression between conditions, we combined the metacells from both conditions and quantified the change in regulatory effect size between conditions for intronic and distal peak-gene pairs that showed a significant association in at least one condition (Figure 2C; Figure 2D; Table S4)(53, 54). We hypothesized that the change in regulatory effect size between conditions could be modeled as an interaction term in the LMM (e.g., modeling gene expression as a function of peak accessibility, condition, and the interaction between condition and accessibility as fixed effects with patient as a random effect). This differential analysis identified 212,781 peak-to-gene associations with significant changes (FDR-adjusted p-value < 1e-04) in regulatory effect size between the cancer and normal conditions (Figure 2E; Table S4). Of these 212,781 differential peak-to-gene associations, 12,954 differential associations had statistically significant effect sizes in the normal condition but insignificant effect sizes in the cancer condition (Figure 2F; Table S4). These differential associations may represent putative regulatory element-target gene pairings specific to the normal condition. Conversely, 188,363 differential associations had statistically significant effect sizes in the cancer condition, but insignificant effect sizes in the normal condition, and thus may reflect cancer-specific putative regulatory element-target gene pairings (Figure 2F; Table S4). The remaining 11,464 differential peak-to-gene associations had statistically significant effect sizes in both conditions, suggesting potential changes in the magnitude and/or direction of the regulatory effect that these putative regulatory elements exert on target gene expression (Figure 2F; Table S4).

To this end, we further classified the differential peak-to-gene associations based on changes in direction between conditions. The differential peak-to-gene associations with significant effect sizes specific to the normal condition were either positive (n=7,665) or negative (n=5,289), suggesting enhancer or silencer-like regulatory relationships that are specific to the normal condition (Figure 2G; Table S4). Similarly, the differential peak-to-gene associations with significant effect sizes specific to the cancer condition were either positive (n=102,489) or negative (n=85,874), suggesting cancer-specific enhancer or silencer-like regulatory relationships in Basal-like BC cells (Figure 2G; Table S4). The remaining differential peak-to-gene associations with significant effects in both conditions were either positive in both conditions (n=3,796), negative in both conditions (n=2,404), or showed changes in direction between conditions (n=5,264). This suggests the potential for regulatory switching events, in which a regulatory element may target the same gene but with opposing regulatory effects depending on the cell state. Overall, most of the differential peak-to-gene associations showed positive regulatory effects that were cancer-specific, indicative of putative enhancer-gene regulation specific to the cancer condition (Figure 2G; Table S4).

To characterize the putative cancer-specific enhancer-regulated genes in Basal-like BC cells, we first screened the cancer-specific peak-to-gene associations for those that involve upregulated genes (FDR-adjusted p-value < 0.05 & Log2FC >= 0.58) in Basal-like BC cells (cancer condition) relative to luminal progenitor cells (normal condition) profiled by scRNA-seq (71-73). This resulted in 7,167 cancer-specific peak-to-gene associations involving upregulated genes and their effect sizes within each condition were visualized in a heatmap (Figure 3A). The heatmap showed a low enrichment of peak-to-gene effect sizes in the normal condition, but a high enrichment of peak-to-gene effect sizes in the cancer condition (Figure 3A). We observed that 84.4% of the 7,167 cancer-specific peak-to-gene associations involved peaks annotated by ENCODE, suggesting they are bona fide enhancers while the remaining likely represent previously unannotated enhancers (Figure 3A)(69, 70). In terms of function, the 7,167 cancer-specific peak-to-gene associations involved 829 unique genes that were enriched (FDR-adjusted p-value < 0.05) for the Hallmark proliferation-associated gene sets E2F TARGETS, G2M CHECKPOINT, and MITOTIC SPINDLE from the Molecular Signatures Database (MSigDB) (Figure 3B)(74-76). Together, these observations provide support for the putative cancer-specific enhancers we identified and suggest that the activities of these enhancers, specifically in Basal-like BC cells, may play critical roles in upregulating genes involved in proliferation.

**Figure 3.**
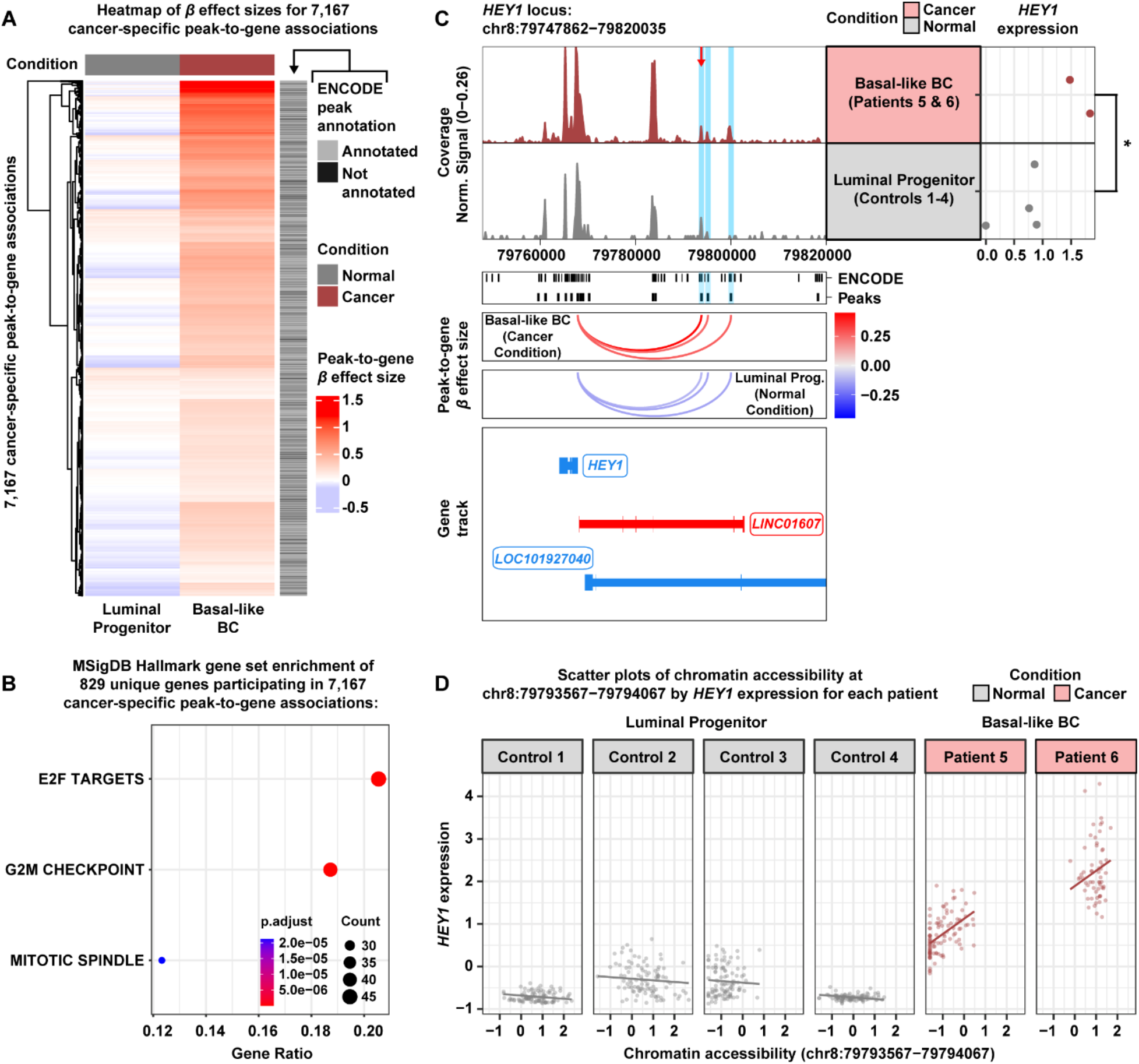
Cancer-specific enhancer regulation of *HEY1* expression in Basal-like BC cells. **A)**. Heatmap of effect sizes for 7,167 cancer-specific peak-to-gene associations in the normal condition, comprised of luminal progenitor cells, and in the cancer condition, comprised of Basal-like BC cells (*left*). Each row represents a peak-gene pair with a significant change in effect size between conditions. The ENCODE peak annotation column denotes the ENCODE annotation status for each cancer-specific peak-to-gene association (*right*). **B).** Hallmark gene set enrichment analysis of 829 unique genes participating in 7,167 cancer-specific peak-to-gene associations as shown in panel **A. C).** Browser track showing the accessibility profile at the *HEY1* locus in the cancer (red) and normal (gray) conditions (*top left*). The putative cancer-specific enhancer with the highest effect size (β = 0.44, FDR-adjusted p-value < 1e-04) on *HEY1* expression is highlighted in light blue and marked by the red arrow. The nearest neighboring putative cancer-specific enhancers (β = 0.25 and β = 0.27, FDR-adjusted p-value < 1e-04) are also highlighted in light blue. Matching pseudo-bulk scRNA-seq expression of *HEY1* is shown for each condition (*top right*). The asterisk denotes a statistically significant difference in gene expression between conditions (FDR-adjusted p-value < 0.05 & Log2FC >= 0.58). ENCODE regulatory element annotations and peaks called from the scATAC-seq data, are shown below the browser track (*middle*). Peak-to-gene loops show the standardized effect sizes, in each condition, of chromatin accessibility at the putative cancer-specific enhancers on *HEY1* expression (*bottom*). **D).** Scatter plots of chromatin accessibility at the strongest putative cancer-specific enhancer by the inferred level of *HEY1* expression in scATAC-seq metacells, stratified by patient in the normal (*gray*) and cancer (*red*) conditions.

We demonstrate a specific example of cancer-specific enhancer-gene regulation in Basal-like BC cells for the upregulated gene *HEY1* (FDR-adjusted p-value < 0.05 & Log2FC >= 0.58) which was linked to 6 putative cancer-specific enhancers (Figure S4; Table S4)(53, 54). *HEY1* is a direct target of the Notch signaling pathway and encodes for Hairy/enhancer-of-split related with YRPW motif protein 1, a basic helix– loop–helix (bHLH) transcription factor (77, 78). Elevated Notch signaling has been observed in a variety of cancers including Basal-like BC (79-84). The cancer-specific enhancer with the highest regulatory effect size (β = 0.44, FDR-adjusted p-value < 1e-04) on *HEY1* expression was annotated by ENCODE, but did not show a statistically significant increase in chromatin accessibility in Basal-like BC cells relative to luminal progenitor cells (Figure 3C) (48, 69-71). This suggests that the putative regulatory element is accessible in both conditions, but targets *HEY1* only in Basal-like BC cells. The nearest neighboring cancer-specific enhancers showed similar regulatory effects on *HEY1* expression (β = 0.25 and β = 0.27, FDR-adjusted p-value < 1e-04), both of which were annotated by ENCODE and the second neighboring cancer-specific enhancer had a statistically significant increase in chromatin accessibility in Basal-like BC cells (Figure 3C) (48, 69-71).

To further visualize the putative regulatory effects of these cancer-specific enhancers on *HEY1* expression, we plotted the levels of chromatin accessibility at the cancer-specific enhancers by the inferred levels of *HEY1* expression in the scATAC-seq metacells from each patient (Figure 3D; Figure S5). This showed that variation in chromatin accessibility at the strongest cancer-specific enhancer was not associated with variation in *HEY1* expression in the luminal progenitor cells from healthy controls, but was significantly associated with variation in *HEY1* expression in the Basal-like BC cells from Patients 5 and 6 (Figure 3C; Figure 3D). The same was observed for the nearest neighboring cancer-specific enhancers (Figure 3C; Figure S5).

Together, these observations suggest a possible mechanism for *HEY1* upregulation in Basal-like BC cells through the activity of tumor unique enhancers that target gene expression in a cancer-specific manner. We note that this example of cancer-specific enhancer-gene regulation is only a glimpse of the altered regulatory landscape in Basal-like BC cells, and we have tabulated all putative regulatory element-target gene pairings identified from the Basal-like subtype analysis in Table S4 that serves as a resource for future investigations of these regulatory elements.

### Cancer-specific regulatory activity of enhancers in Luminal BC cells

Luminal BC is often associated with hormone receptor positive BC, and is the most commonly diagnosed BC among women(62, 85). To analyze the cells in the Luminal subtype analysis, we merged Luminal BC cells from Patients 7-15 with mature luminal cells from the healthy mammary reduction patients, according to the results of the unsupervised pseudo-bulk clustering analysis (Figure S3). This subset resulted in 13,351 cells and 15,883 cells profiled by scRNA-seq and scATAC-seq, respectively (Figure 4A; Table S2C; Table S3C). After re-clustering the scRNA-seq cells and transferring the resulting labels as well as gene expression profiles to the scATAC-seq cells, we observed that Luminal BC cells mainly clustered by patient, consistent with previous reports, and the mature luminal cells from healthy controls were represented by a single cluster (Figure 4B; Table S2C; Table S3C)(23, 28, 58-60).

**Figure 4.**
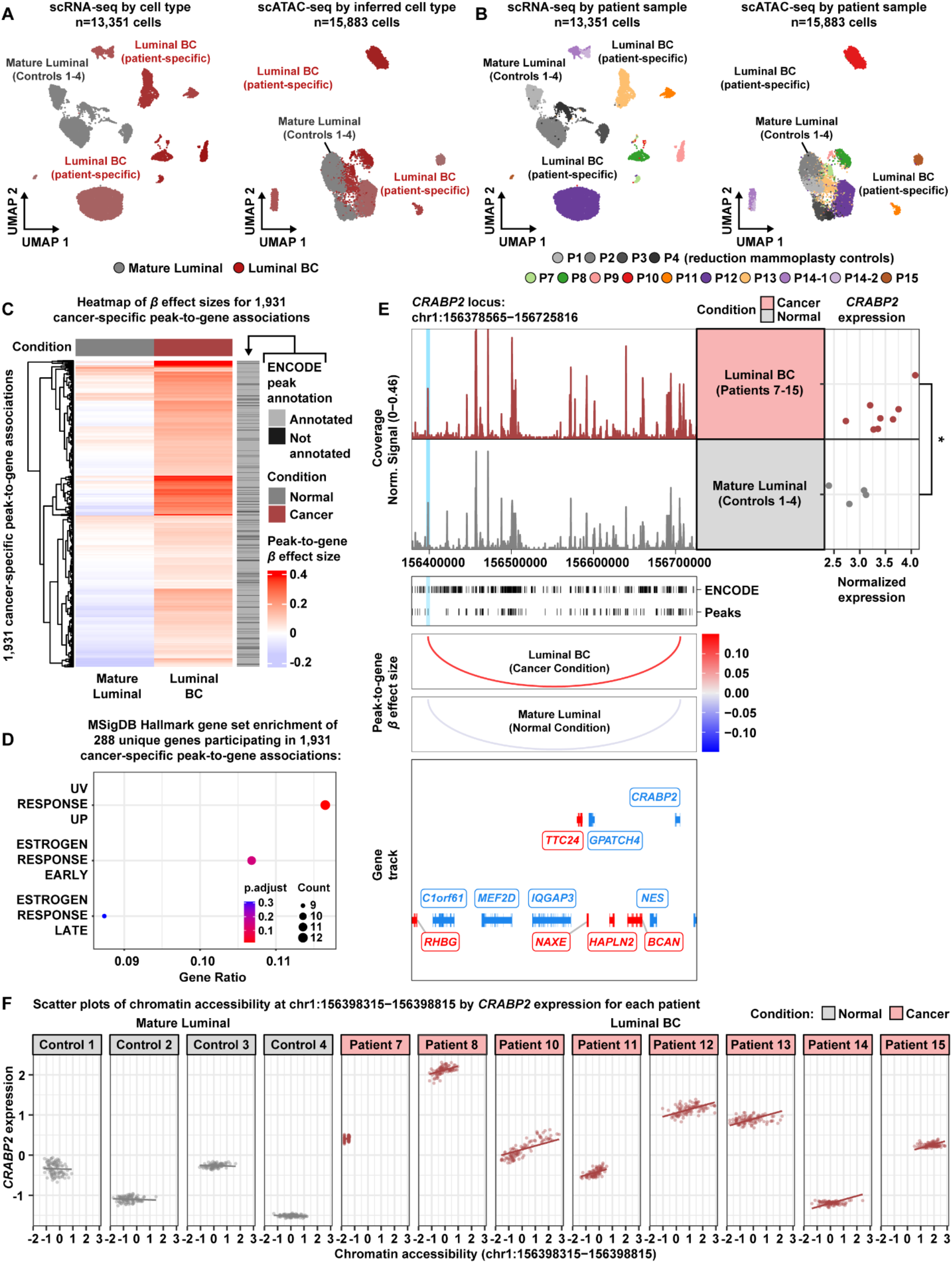
Cancer-specific enhancer regulation of *CRABP2* expression in Luminal BC cells. **A).** UMAP plot of 13,351 scRNA-seq cells color-coded by cell type across 14 patient samples (*left*). UMAP plot of 15,883 scATAC-seq cells color-coded by inferred cell type across 14 patient samples (*right*). Color shades denote clusters within each cell type**. B).** UMAP plots of scRNA-seq cells (left) and scATAC-seq cells (right), as shown in panel **A**, but color-coded by the patient sample of origin. **C).** Heatmap of effect sizes for 1,931 cancer-specific peak-to-gene associations in the normal condition, comprised of mature luminal cells, and in the cancer condition, comprised of Luminal BC cells (*left*). Each row represents a peak-gene pair with a significant change in effect size between conditions. ENCODE peak annotation column denotes the ENCODE annotation status for each cancer-specific peak-to-gene association (*right*). **D).** Hallmark gene set enrichment analysis of 288 unique genes participating in 1,931 cancer-specific peak-to-gene associations, as shown in panel **C. E).** Browser track showing the accessibility profile at the *CRABP2* locus for the cancer (red) and normal (gray) conditions (*top left*). The putative cancer-specific enhancer with the highest effect size (β = 0.11, FDR-adjusted p-value < 1e-04) on *CRABP2* expression is highlighted in light blue. Matching pseudo-bulk scRNA-seq expression of *CRABP2* is shown for each condition (*top right*). The asterisk denotes a statistically significant difference in gene expression between conditions (FDR-adjusted p-value < 0.05 & Log2FC >= 0.58). ENCODE regulatory element annotations and peaks called from the scATAC-seq data, are shown below the browser track (*middle*). Peak-to-gene loops show the standardized effect size, in each condition, of chromatin accessibility at the putative cancer-specific enhancer on *CRABP2* expression (*bottom*). **F).** Scatter plots of chromatin accessibility at the putative cancer-specific enhancer by the inferred level of *CRABP2* expression in scATAC-seq metacells, stratified by patient in the normal (*gray*) and cancer (*red*) conditions.

To probe the altered regulatory landscape in Luminal BC cells, relative to mature luminal cells, we carried out peak calling and performed the two-phased differential peak-to-gene association framework as performed in the Basal-like subtype analysis (Figure 2C; Figure S6)(13, 48, 53, 54, 67, 68). The first phase of the differential peak-to-gene association analysis, quantifying peak-to-gene regulatory effect sizes within each condition independently, yielded similar results to the Basal-like subtype analysis with a total of 430,119 significant peak-to-gene associations (FDR-adjusted p-value < 1e-04), most of which involved peaks annotated by the ENCODE database and were located in introns and distal intergenic regions (Figure S6A; Table S5) (23, 69, 70).

The second phase of the differential peak-to-gene association analysis, quantifying changes in peak-to-gene regulatory effect size between conditions, also yielded similar results to the Basal-like subtype analysis with a total of 135,633 significant differential peak-to-gene associations (FDR-adjusted p-value < 1e-04). We note that 5,859 differential associations showed significant changes in direction which again may be interpreted as possible regulatory switching events. However, most the differential peak-to-gene associations showed positive effect sizes specific to the cancer condition of Luminal BC cells, which again may be interpreted as putative enhancers targeting gene expression in a cancer-specific manner (Figure S6B-D; Table S5).

There were 1,931 cancer-specific peak-to-gene associations involving upregulated genes (FDR-adjusted p-value < 0.05 & Log2FC >= 0.58) and their effect sizes within each condition were visualized in a heatmap (Figure 4C)(71-73). 91.2% of the 1,931 cancer-specific peak-to-gene associations involved peaks annotated by ENCODE (Figure 4C)(69, 70). To assess function, we observed that the 1,931 cancer-specific peak-to-gene associations involved 288 unique genes that were enriched (FDR-adjusted p-value < 0.05) for the Hallmark DNA damage-associated gene set UV RESPONSE UP, from MSigDB (Figure 4D)(74-76). We also observed that the next most significant enrichments were for the Hallmark signaling-associated gene sets: ESTROGEN RESPONSE EARLY and ESTROGEN RESPONSE LATE, however these gene sets did not reach statistical significance (FDR-adjusted p-value > 0.05)(Figure 4D). Together, these observations suggest that the activity of these putative cancer-specific enhancers in Luminal BC cells may offer mechanistic insights into the regulation of DNA damage repair pathways in response to estrogen(86, 87).

We highlight a specific example of cancer-specific enhancer-gene regulation in Luminal BC cells for the upregulated gene *CRABP2* (FDR-adjusted p-value < 0.05 & Log2FC >= 0.58), which was linked to 14 putative cancer-specific enhancers (Figure S7; Table S5)(53, 54). *CRABP2* encodes for cellular retinoic acid binding protein 2 that shuttles retinoic acid from the cytosol to the nucleus (88, 89). Interestingly, high expression of *CRABP2* has been reported in a number of cancers, including BC(90-99). The cancer-specific enhancer with the highest regulatory effect size (β = 0.11, FDR-adjusted p-value < 1e-04) on *CRABP2* expression was annotated by ENCODE, but did not show a statistically significant increase in chromatin accessibility in Luminal BC cells (Figure 4E)(48, 69-71). This suggests that the putative regulatory element is accessible in both conditions, but targets *CRABP2* only in Luminal BC cells.

To visualize this proposed mechanism, we plotted the levels of chromatin accessibility at the cancer-specific enhancer by the inferred levels of *CRABP2* expression in the scATAC-seq metacells from each patient (Figure 4F). We observed that variation in chromatin accessibility at this cancer-specific enhancer was not associated with variation in *CRABP2* expression in the mature luminal cells from healthy controls but was significantly associated with variation in *CRABP2* expression in the Luminal BC cells from each BC patient.

Together, these observations describe a potential mechanism for *CRABP2* upregulation in Luminal BC cells. We note that this specific example of cancer-specific enhancer-gene regulation is only a snap shot of the altered regulatory landscape in Luminal BC cells and the remaining putative regulatory element-target gene associations are tabulated in Table S5.

### Clinical relevance and essentiality of cancer-specific enhancer-regulated genes

To evaluate the clinical relevance of *HEY1* and *CRABP2* in subtype-specific BCs, we performed survival analyses and assessed their copy number variation (CNV) status in external patient cohorts. We fit a Cox proportional hazards model for *HEY1* using TNBC patients in each dataset (Methods). The same was performed for *CRABP2* using HR+/HER2-patients in each dataset. These analyses revealed that high expression of *HEY1* was associated with worse outcome in three out of five TNBC patient datasets (HR >1, Cox p-value < 0.01) while high expression of *CRABP2* was associated with worse outcome in two out of three HR+/HER2-patient datasets (HR >1, Cox p-value < 0.01), respectively (Table S6). We further visualized these trends using Kaplan-Meier plots with log-rank p-values displayed (HR >1, log-rank p-value < 0.01) (Figure 5A-B). These analyses provide evidence to suggest that *HEY1* and *CRABP2* are prognostic in TNBC and HR+/HER2-patient populations, respectively. We next visualized the frequencies of CNV that affect *HEY1* and *CRABP2* in TCGA-BRCA patients. This revealed a majority of TNBC patients that show copy number gains near the *HEY1* locus on chromosome 8 (Figure 5C). The same was observed for the *CRABP2* locus on chromosome 1 in HR+/HER2-patients (Figure 5D). These observations suggest that *HEY1* and *CRABP2* may be co-amplified with their putative cancer-specific enhancers, serving as additional mechanisms of upregulation.

**Figure 5.**
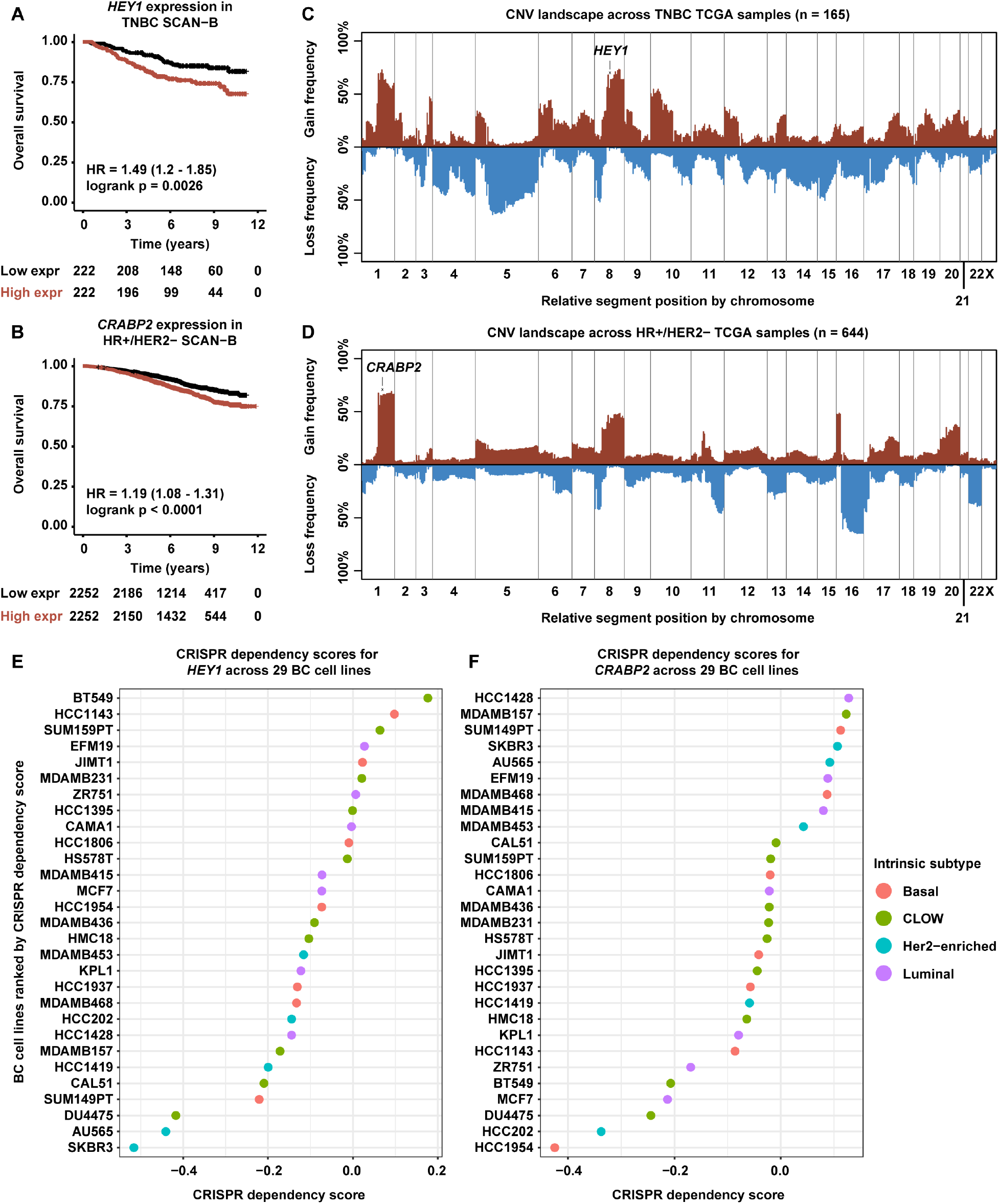
Clinical and biological relevance of cancer-specific enhancer-regulated genes. **A).** Kaplan-Meier survival curve based on event-free survival for 444 TNBC SCAN-B patients stratified by high and low *HEY1* expression. **B).** Kaplan-Meier survival curve based on event-free survival for 4504 HR+/HER2-SCAN-B patients stratified by high and low *CRABP2* expression. **C).** CNV landscape plot showing copy number gain or loss frequencies in 165 TNBC TCGA patients by chromosomal segment. The *HEY1* locus is labeled with an x. **D).** CNV landscape plot, as in panel **C**, but for 644 HR+/HER2-TCGA patients. The *CRABP2* locus is labeled with an x. **E).** Dot plot of *HEY1* CRISPR dependency scores by BC cell line color coded by intrinsic subtype including claudin-low (CLOW). **F).** Dot plot, as in panel **E**, but for *CRABP2* CRISPR dependency scores.

To evaluate the essentiality of *HEY1* and *CRABP2* in subtype-specific BCs, we analyzed CRISPR dependency scores in subtype-specific BC cell lines from the DepMap portal (100). CRISPR dependency scores quantify the effect of knocking out a target gene on cell growth and proliferation of a particular cell line. We visualized CRISPR dependency scores for *HEY1* in 29 subtype-specific BC cell lines. Interestingly, the majority of Basal-like cell lines showed dependency scores less than zero indicating a growth deficit after *HEY1* knockout and the Basal-like cell line SUM149PT was the most sensitive (Figure 5E). To confirm that *HEY1* is expressed in Basal-like cell lines from the Cancer Cell Line Encyclopedia (CCLE), we visualized the distribution of *HEY1* expression (RNA-seq) across all 29 BC cell lines stratified by subtype, revealing modest expression across all subtypes (Kruskal-Wallis rank sum test, p-value > 0.05) (Figure S8A)(101). Similarly, we visualized CRISPR dependency scores for *CRABP2* in the same set of BC cell lines. While the top four most sensitive cell lines to *CRABP2* knockout only included one Luminal cell line (MCF7), we note that the majority of Luminal cell lines were sensitive to *CRABP2* knockout (Figure 5F). Moreover, we also observed that *CRABP2* is highly expressed in Luminal cell lines (Kruskal-Wallis rank sum test, p-value < 0.05) (Figure S8B). Together, these observations suggest that cancer-specific enhancer-regulated genes may play essential roles in the growth and proliferation of BC cells *in vitro*.

### Annotation of the enhancer regulatory landscapes in subtype-specific BC cells *in vitro*

To investigate enhancer-regulated gene expression in subtype-specific BC cells *in vitro*, we also generated matched scRNA-seq and scATAC-seq profiles for the Basal-like BC cell lines HCC1143 and SUM149PT, and the Luminal BC cell lines MCF7 and T47D (Figure 1A; Table 1)(102). We carried out QC, dimensionality reduction, and cross-modality integration in the cell line dataset as performed in the patient datasets. This resulted in 30,651 cells and 16,338 cells profiled by scRNA-seq and scATAC-seq, respectively (Table 1; Table S2D; Table S3D; Figure 6A-B; Figure S9).

**Figure 6.**
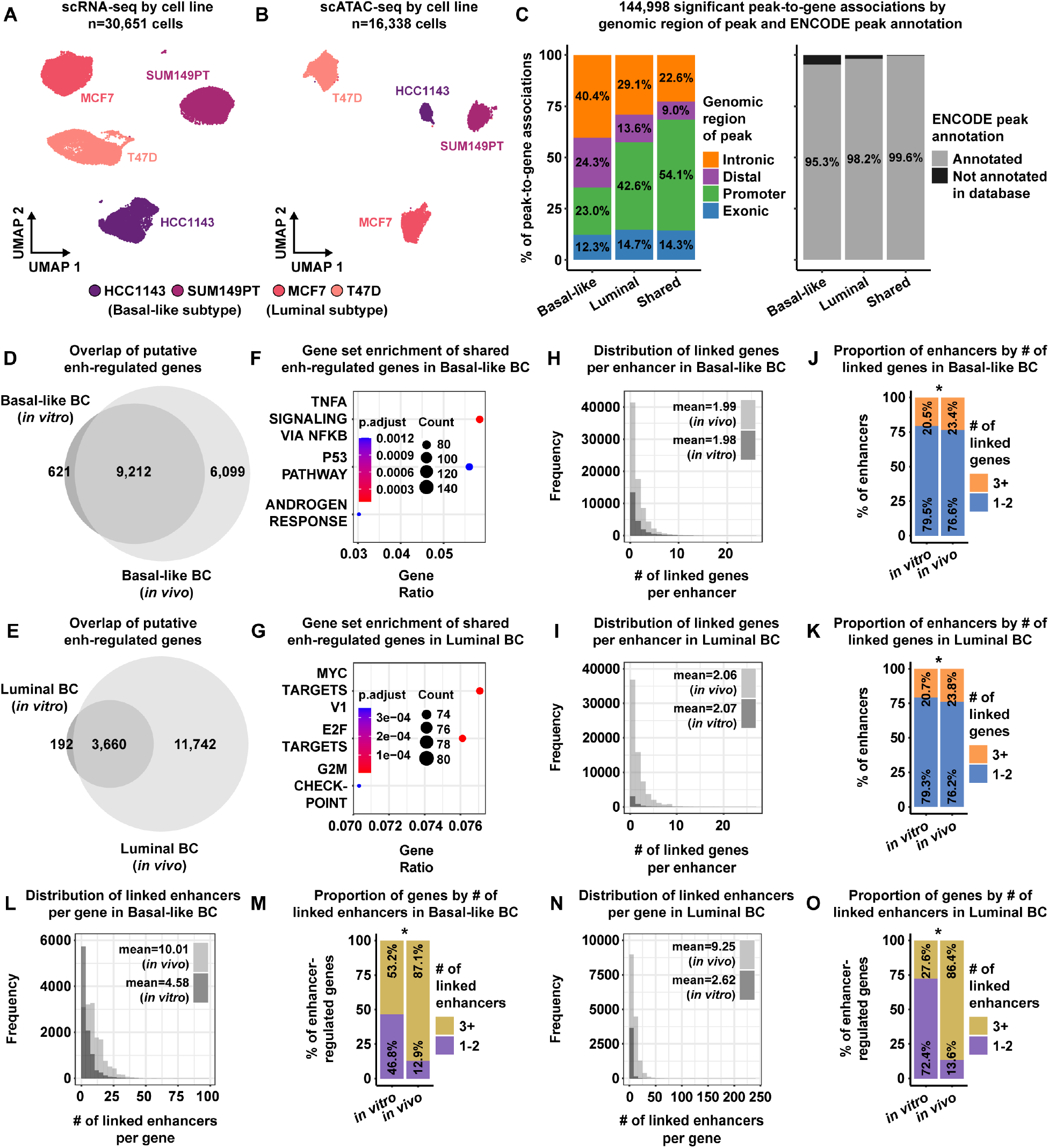
Comparison of enhancer regulatory landscapes between *in vitro* and *in vivo* subtype-specific BC cells. **A)**. UMAP plot of 30,651 scRNA-seq cells color-coded by cell line across 4 cell line samples. **B)**. UMAP plot of 16,338 scATAC-seq cells color-coded by inferred cell line across 4 cell line samples. **C)**. Proportion bar charts showing the genomic distribution (*left*) and ENCODE annotation status (*right*) for 105,884 Basal-like-specific peak-to-gene associations, 30,998 Luminal-specific associations, and 8,116 shared associations. **D)**. Venn diagram showing the overlap of putative enhancer-regulated genes between Basal-like BC cells *in vitro* and *in vivo*. **E)**. Venn diagram showing the overlap of putative enhancer-regulated genes between Luminal BC cells *in vitro* and *in vivo*. **F)**. Hallmark gene set enrichment analysis of 9,212 shared enhancer-regulated genes between Basal-like BC cells *in vitro* and *in vivo*. **G)**. Hallmark gene set enrichment analysis of 3,660 shared enhancer-regulated genes between Luminal BC cells *in vitro* and *in vivo*. **H)**. Histograms showing the distributions of linked genes per enhancer for Basal-like BC cells *in vitro* and *in vivo*. **I)**. Histograms, as in panel **H**, but for Luminal BC cells *in vitro* and *in vivo*. **J)**. Proportion bar charts showing the proportions of enhancers by number of linked genes for Basal-like BC cells *in vitro* and *in vivo*. The asterisk denotes a statistically significant difference in the proportion of enhancers that link to three or more genes between Basal-like BC cells *in vitro* and *in vivo*. **K)**. Proportion charts, as in panel **J**, but for Luminal BC cells *in vitro* and *in vivo*. The asterisk denotes a statistically significant difference in the proportion of enhancers that link to three or more genes between Luminal BC cells *in vitro* and *in vivo*. **L)**. Histograms showing the distributions of linked enhancers per gene for Basal-like BC cells *in vitro* and *in vivo*. **M)**. Proportion bar charts showing the proportions of genes by number of linked enhancers for Basal-like BC cells *in vitro* and *in vivo*. The asterisk denotes a statistically significant difference in the proportion of genes that link to three or more enhancers between Basal-like BC cells *in vitro* and *in vivo*. **N)**. Histograms, as in panel **L**, but for Luminal BC cells *in vitro* and *in vivo*. **O)**. Proportion bar charts, as in panel **M**, but for Luminal BC cells *in vitro* and *in vivo*. Asterisk denotes a statistically significant difference in the proportion of genes that link to three or more enhancers between Luminal BC cells *in vitro* and *in vivo*.

To link putative regulatory elements to target genes in subtype-specific BC cells *in vitro*, we carried out peak calling in scATAC-seq cells and quantified peak-to-gene regulatory effect sizes, using our robust LMM-based approach, within each subtype independently (13, 48, 53, 54, 67, 68). This revealed 144,998 significant peak-to-gene associations, with 105,884 associations specific to Basal-like BC cells *in vitro*, 30,998 associations specific to Luminal BC cells *in vitro*, and 8,116 shared associations (Figure 6C; Table S7). We observed that the majority of the Basal-like-specific peak-to-gene associations involved peaks located in introns and distal intergenic regions, while the majorities of Luminal-specific and shared associations involved peaks located in promoters and exonic regions (Figure 6C; Table S7). Again, we observed that a strong majority of these peak-to-gene associations involved peaks annotated by ENCODE (Figure 6C; Table S7)(69, 70).

We next sought to compare the enhancer regulatory landscapes between BC cells *in vitro* and *in vivo* stratified by molecular subtypes (e.g., comparing *in vitro* Basal-like BC cells with *in vivo* Basal-like BC cells). To this end, we screened the peak-to-gene associations identified in subtype-specific BC cells *in vitro* for those with positive effect sizes. The same screening was done for the peak-to-gene associations identified *in vivo* from the subtype-specific patient analyses (Figure 2C-D; Figure S6A). We then performed overlap analyses of putative enhancer-regulated genes between *in vitro* and *in vivo* subtype-specific BC cells (Figure 6D-E; Methods).

94% and 95% of enhancer-regulated genes *in vitro* were also enhancer-regulated *in vivo*, for Basal-like and Luminal BC cells, respectively (Figure 6D-E). These shared enhancer-regulated genes in Basal-like BC cells were enriched (FDR-adjusted p-value < 0.05) for the Hallmark signaling-associated gene sets TNFA SIGNALING VIA NFKB and ANDROGEN RESPONSE as well as the Hallmark proliferation-associated gene set P53 PATHWAY from MSigDB (Figure 6F). In Luminal BC cells, shared enhancer-regulated genes were enriched (FDR-adjusted p-value < 0.05) for the Hallmark proliferation-associated gene sets: MYC TARGETS V1, E2F TARGETS, and G2M CHECKPOINT (Figure 6G).

To quantify the “regulatory load” of the putative enhancers identified in each setting, we visualized the distributions of linked genes per enhancer for the enhancers identified *in vitro* and *in vivo*, stratified by molecular subtype (Figure 6H-I). In Basal-like BC cells, the mean numbers of linked genes per enhancer *in vitro* and *in vivo* were 1.98 and 1.99, respectively (Figure 6H). In Luminal BC cells, the mean numbers of linked genes per enhancer *in vitro* and *in vivo* were 2.07 and 2.06, respectively (Figure 6I). We also observed that 23.4% of enhancers identified in Basal-like BC cells *in vivo* linked to 3 or more genes, compared to only 20.5% of enhancers identified *in vitro* (OR=1.19, p-value < 0.01, Fisher’s exact test) (Figure 6J). In Luminal BC cells, we observed that 23.8% of enhancers identified *in vivo* linked to 3 or more genes, compared to only 20.7% of enhancers identified *in vitro* (OR=1.2, p-value < 0.01, Fisher’s exact test) (Figure 6K). The same analyses were performed for the number of linked enhancers per gene (Figure 6L-O).

In Basal-like BC cells, we observed that the mean numbers of linked enhancers per gene *in vitro* and *in vivo* were 4.58 and 10.01, respectively (Figure 6L). 87.1% of enhancer-regulated genes in Basal-like BC cells *in vivo* linked to 3 or more enhancers, compared to only 53.2% of enhancer-regulated genes *in vitro* (OR=5.92, p-value < 0.01, Fisher’s exact test) (Figure 6M). Similarly, we observed that the mean numbers of linked enhancers per gene in Luminal BC cells *in vitro* and *in vivo* were 2.62 and 9.25, respectively (Figure 6N). 86.4% of enhancer-regulated genes in Luminal BC cells *in vivo* linked to 3 or more enhancers, compared to only 27.6% of enhancer-regulated genes *in vitro* (OR=16.69, p-value < 0.01, Fisher’s exact test) (Figure 6O).

Overall, these observations suggest that enhancers *in vivo* may regulate more genes compared to enhancers *in vitro* and that genes expressed *in vivo* may be regulated by more enhancers compared to genes expressed *in vitro*, possibly due to the clonal heterogeneity and/or tumor microenvironment factors that BC cells experience *in vivo*.

## DISCUSSION

BC is the most commonly diagnosed cancer among women and accounts for a significant proportion of female cancer-related deaths, highlighting the need for deeper insights into the molecular underpinnings of BCs that may lead to improved targeted therapies(1). The compendium presented herein represents a valuable multi-omic resource that unveils the transcriptional and regulatory landscapes of human breast tumors and normal mammary epithelial tissues at single-cell resolution. More specifically, our work elucidated the transcriptional and regulatory features that distinguish BC cells from their nearest normal precursor cell types by identifying putative enhancers that regulate clinically relevant oncogenic expression programs in a cancer-specific manner (Figures 2-6; Figures S3-S7; Tables S4-5) (31, 32, 35, 53, 54, 71, 74-76, 103). These data also enabled us to study the transcriptional and regulatory differences between BC cells *in vitro* and *in vivo* (Figure 6; Table S7)(53, 54, 74-76).

We reiterate three important themes from analyzing these single-cell data. First, we demonstrated how our computational approach, for linking putative regulatory elements to target genes, accounts for important biological and technical variables when quantifying associations between chromatin accessibility and gene expression (Figures 2-4; Figure 6; Figures S4-S7; Tables S4-5, Table S7) (53, 54). It has become widely accepted that the activity of *cis*-regulatory elements is highly cell type-specific, therefore, it is critical to stratify by cell type when quantifying peak-to-gene associations from single-cell multi-omic data (Figure 2C) (43-45). Accounting for potential batch and/or sample-specific effects is also important to ensure that technical variation in chromatin accessibility and/or gene expression is not confused with biological variation relevant to the hypothesis (104, 105).

Our computational approach also provides a measure of statistical significance for changes in peak-to-gene regulatory effect size between conditions and/or cell types (53, 54). This allows for inferences about possible changes in the magnitude and/or direction of the regulatory effect that a regulatory element exerts on target gene expression. Our differential peak-to-gene association analysis allowed us to classify peak-to-gene associations based on changes in direction of effect size between conditions. For both the Basal-like and Luminal subtype analyses, this revealed thousands of cancer-specific associations with positive effect sizes indicative of context-specific putative enhancer activity and evidence to suggest the potential for regulatory switching events that were previously hidden using current peak-to-gene association methods (Figure 2E-G; Figure S6B-D) (48, 106-108).

Next, we highlight that the putative cancer-specific enhancers, identified in Basal-like and Luminal BC cells, were linked to genes involved in known oncogenic processes including proliferation and DNA damage, respectively (Figure 3B; Figure 4D)(74-76). Moreover, we were able to show specific examples of cancer-specific enhancer regulation that not only may be associated with clinical outcomes, but also may be amplified through copy number alterations (Figure 5A-D; Table S6)(103). We also observed that these cancer-specific enhancers linked to genes that may be essential for BC cell proliferation and survival *in vitro*, suggesting there may exist upstream “regulatory dependencies” that modulate the expression of essential genes (Figure 5E-F). Together, these observations further underscore the importance of non-coding regulatory mechanisms for transcriptional dysregulation in BC cells.

Finally, in the comparisons of subtype-specific BC cells *in vitro* to *in vivo*, our analyses point to a conserved set of genes expressed in both settings for each subtype. We note that these conserved sets of expressed genes are associated with known oncogenic processes relevant to BC cells both *in vitro* and *in vivo*, including TNF-alpha signaling and proliferation (Figure 6D-G)(74-76, 109-112). We speculate that perhaps enhancer rewiring or hijacking may be a stochastic process that gets selected for when a specific set of genes, favorable to the cancer cells, become upregulated. In summary, this resource demonstrates important principles of enhancer-gene regulation in BC cells profiled by single-cell multi-omics and serves as an important resource to the field.

### Limitations of study

We acknowledge that there are some limitations to our study. First, the libraries for scRNA-seq and scATAC-seq were derived from separate, albeit homogeneous, aliquots of cell suspensions for each patient specimen or cell line. This experimental design requires the downstream use of statistical tools for cross-modality integration of these single-cell data, unlike recent methods for profiling the transcriptional and chromatin landscape within the same cell(35, 106). However, the cell recovery rate and sequencing depth of these “same cell” protocols can be lower compared to scRNA-seq and scATAC-seq, which can affect the accuracy of downstream analyses (113). We were also able to validate the performance of the cross-modality integration method used in our study by leveraging the ground-truth cell identities of scATAC-seq cells in the cell line data. Second, we recognize that our survival analyses involved gene expression measurements derived from bulk tissues in contrast to these single-cell data. Third, we realize that our comparisons of BC cells from BC patients to normal mammary epithelial cells from healthy controls may be limited by possible confounding biological factors such as age and menopause status. However, we note that our normal control cells from healthy controls represent a true baseline for gene regulation in mammary epithelial tissues, unlike tumor adjacent normal tissues (used in many studies) that may be affected by tumor microenvironment factors and genomic alterations (114-116). Finally, we acknowledge that our study had a limited sample size, especially for patient tumors of the TNBC clinical and/or Basal-like molecular subtypes, which could affect the generalizability of the resource and our observations. However, our study focused on treatment-naïve breast tumors which are difficult to procure as the standard of care shifts towards neoadjuvant treatment prior to surgery(117-120). Nonetheless, the resource and our analyses described herein provide an unobscured view of gene regulation in BC cells and highlights potential avenues for therapeutic interventions.

## METHODS

### Human patient samples and tissue dissociation

Eleven, treatment naïve, breast cancer patients were enrolled in the 2018 Breast SPORE Project 2 study at the UNC Cancer Hospital (IRB Protocol 17-3228) and underwent curative intent surgical resection (Table 1; Table S1). Additionally, four patients were enrolled who were undergoing reduction mammoplasty surgeries in order to collect normal control samples (Table 1; Table S1). After surgical resection, tissue specimens were sectioned by the pathology department and the remaining tissues were de-identified and collected for this study through the University of North Carolina’s Tissue Procurement Facility. The tissue specimens were never fixed or frozen and were transported to the lab immediately after surgical resection on ice in media containing DMEM/F12 media (Gibco) + 1% Penicillin/Streptomycin (Corning). Tissues were dissociated as previously reported(23). In short, before dissociation, tumor and normal samples were weighed. Tissue mass varied between 0.12 g and 11.5 g. Tissue specimens were then minced and digested overnight in DMEM/F12 + 5% FBS, 15mM HEPES (Gibco), 1x Glutamax (Gibco), 1x Collagenase/Hyaluronidase (Stem Cell Technologies, 07912), 1% Penicillin/Streptomycin (Corning), and 0.48 µg/mL Hydrocortisone (Stem Cell Technologies, 74144) on a stir plate at 37□C and 180 rpm. Some tissues were dissociated with Gentle Collagenase/Hyaluronidase (Stem Cell Technologies, 07919) instead of Collagenase/Hyaluronidase. After digestion, cells were washed twice with cold PBS + 2% FBS and 10mM HEPES (PBS-HF) and centrifuged. To remove red blood cells, the cell pellet was treated with cold Ammonium Chloride Solution (Stem Cell Technologies, 07850) and PBS-HF (ratio 1 Ammonium Chloride: 4 PBS-HF), for 1 minute, then centrifuged. The volume of Ammonium Chloride Solution added was determined by the size of the cell pellet and visual assessment of pink or red color of the pellet. Red blood cell removal was repeated a second time if the pellet still exhibited a pink color after the first treatment. Next, cell pellets were resuspended in 0.05% Trypsin-EDTA (Gibco) and the suspension was gently pipetted up and down for 1 min. Trypsin was then inactivated by adding 10mL PBS-HF solution and the suspension was centrifuged. If cell suspensions were still clumpy after trypsin treatment, cells were resuspended with 1-2 mL Dispase (Stem Cell Technologies, 07923) and 200 µL 1mg/mL DNase I (Stem Cell Technologies, 07900) for 1 min, then inactivated with 10 mL PBS-HF. If the Dispase step was not necessary, cells were treated with DNase I during the trypsinization step. Cells were again centrifuged, then washed in PBS-HF and filtered through a 100µm cell strainer and washed again. The cell pellet was resuspended in DMEM/F12 + 5% FBS using a volume based on the final pellet size and filtered using a 40µm cell strainer. Single-cell suspension concentration and cell viability was measured with the Countess II Automated Cell Counter (Thermo Fisher, AMQAX1000). Cell viability varied between 43% and 94% across all samples, with the majority of suspensions having over 70% viability.

### Cell culture

Cell lines were obtained from the American Type Culture Collection (ATCC) and maintained in the Franco lab at UNC Chapel Hill. The MCF-7 and T47D cells were grown in Dulbecco’s Modified Eagle Medium (DMEM) (Sigma cat. #30-2002) supplemented with 10% fetal bovine serum (FBS) (Sigma) and 1% penicillin/streptomycin (Corning). The HCC1143, and SUM149PT cells were grown in RPMI-1640 media (Sigma) supplemented with 10% FBS (Sigma) and 1% penicillin/streptomycin (Corning). All cell lines were grown adherent in a 5% CO2 incubator set to 37°C with low passage stocks reserved in liquid nitrogen. Cell lines were authenticated by ATCC and tested for mycoplasma prior to use.

### Single-cell sequencing

Cell suspensions were next used for scRNA-seq and scATAC-seq library prep. For scRNA-seq, cell suspensions were diluted to 1200 cells/µL and 10,000 cells were used in library generation with the following 10x Genomics Single Cell 3’ kits: Chromium Single Cell 3’ GEM, Library & Gel Bead Kit v3 (PN-1000075), Chromium Chip B Single Cell Kit (PN-10000153), and Chromium i7 Multiplex Kit (PN-120262) following the manufacturer’s protocol.

For scATAC-seq, 500,000 cells were used in nuclei isolation following the Nuclei Isolation for Single Cell ATAC Sequencing protocol from 10x Genomics. For the lysis step, cells were lysed for 4 min. For the resuspension step, nuclei were resuspended in 50 µL 1x Nuclei Buffer. Nuclei were counted with the Countess II Automated Cell Counter. 10,000 nuclei were used in library preparation using the following 10x Genomics Single Cell ATAC Kits: Chromium Single Cell ATAC Library & Gel Bead Kit v1 (PN-1000110), Chromium Chip E Single Cell ATAC Kit (PN-1000082), and Chromium i7 Multiplex Kit N, Set A (PN-1000084) following the manufacturer’s protocol. All libraries were sequenced using the 10X Genomics suggested sequencing parameters on an Illumina NextSeq 500 instrument.

### Quantification and quality control (QC) in single-cell RNA-seq

Filtered feature barcode matrices were generated for each patient and cell line sample using Cell Ranger (version 3.1.0) from 10x Genomics. For each sample, the filtered feature barcode matrix was converted into a Seurat object using the *CreateSeuratObject()* function from the Seurat R package(46, 47, 121). QC and doublet removal were carried out for each sample individually. Barcodes with at least 500 expressed genes, at least 1,000 UMI counts, and less than 20% mitochondrial counts were deemed high quality cells and were carried forward to doublet detection(122). Cells predicted as doublets by the DoubletFinder R package were removed from further downstream analyses (121, 123). After QC and doublet removal for each sample, Seurat’s *merge()* function was used to concatenate the individual patient samples and the individual cell line samples, to form the patient cohort dataset and the cell line cohort dataset, respectively (46, 47).

### Single-cell RNA-seq normalization, feature selection, and clustering

All gene expression matrices (for individual samples as well as cohort datasets) were normalized with Seurat’s *NormalizeData()* function (46, 47). Seurat’s *FindVariableFeatures()* function was used to identify the top 2,000 most variably expressed genes within each individual sample and within each of the cohort datasets (patient cohort, Basal-like subtype cohort, Luminal subtype cohort, and cell line cohort datasets introduced in Figures 1, 2, 4, and 6, respectively) (46, 47). The data were scaled with Seurat’s *ScaleData()* function using all genes for each individual sample and using only the top variably expressed genes for the remaining cohort datasets(46, 47). For all analyses of each sample individually and each cohort, the top 2,000 most variably expressed genes were used for principal component analysis (PCA) with Seurat’s *RunPCA()* function and cells were visualized in a uniform manifold approximation and projection (UMAP) plot using Seurat’s *RunUMAP()* function with the first 30 principal components (PCs)(46, 47). UMAP plots were then plotted using the ggplot2 R package(121, 124).

For each individual patient sample, cells were clustered using the R package MultiK to identify an optimal number of clusters(121, 125). Note that MultiK applies Seurat’s clustering methods over multiple resolution parameters(46, 47, 125). Cells within each cohort dataset (patient cohort, Basal-like subtype cohort, Luminal subtype cohort, and cell line cohort) were clustered using Seurat’s clustering methods which included building a graph with the *FindNeighbors()* function using the first 30 PCs and identifying clusters with the *FindClusters()* function(46, 47). The resolution parameter in *FindClusters()* was set to 0.4, 0.015, and 0.015 for the patient, Basal-like, and Luminal cohort datasets, respectively.

### Cell type annotation in single-cell RNA-seq

Cell type annotation was initially performed within each individual patient sample after pre-processing and clustering with MultiK(46, 47, 123, 125). For each patient sample, cells were annotated to known cell types using Seurat’s canonical correlation analysis (CCA)-based label transfer procedure with a large scRNA-seq breast cancer (BC) reference dataset downloaded from GSE176078 (28, 46, 47). Clusters within each patient sample were then annotated based on their majority predicted label from the reference-based label transfer procedure. Since mast cells were underrepresented in the reference scRNA-seq dataset, some clusters within each patient dataset were re-annotated to mast cells if the cluster showed significant marker gene expression of *TPSB2* and *TPSAB1* (Bonferroni-corrected p-value < 0.01, log2FC > 0.25) (28, 126). Within each patient sample, cluster annotations were verified by visualizing the distributions of cell type gene signature enrichment scores per cluster using Seurat’s *AddModuleScore()* function with relevant signatures sourced from PanglaoDB (46, 47, 127). The clusters identified in the patient cohort dataset, after combining the individual patient samples, were annotated based on their majority cell type label derived from the annotated clusters in each individual patient sample (Figure 1B-D; Figure S2A-B).

### Inference of copy number variation (CNV), cancer cell identification, and molecular subtype prediction from single-cell RNA-seq

Inferred CNV scores were estimated for individual cells annotated as epithelial within each BC patient sample using the R package inferCNV(28, 55-57). To identify high-confidence cancer cells within each BC patient sample, cells annotated as epithelial were classified into one of three groups: inferCNV high, ambiguous, or inferCNV low, based on the inferred CNV score of each cell as described previously (Figure 1D; Figure S2B) (28, 55-57). Cells classified as inferCNV high were deemed putative cancer cells and were carried forward to molecular subtype prediction (Basal, Her2-enriched, Luminal A, or Luminal B) using the SCSubtype method described previously (Figure 1D; Figure S2B) (28). Briefly, this procedure calculates subtype-specific signature enrichment scores for individual cells assigned to one of four molecular subtypes (Basal, Her2-enriched, Luminal A, or Luminal B) based on the highest signature enrichment score for each cell (28).

### Quality control (QC) in single-cell ATAC-seq

A list of unique ATAC-seq fragments with associated barcodes was generated for each patient and cell line sample using Cell Ranger ATAC (version 1.2.0) from 10x Genomics. These lists of unique ATAC-seq fragments per barcode were read into the ArchR R package using the *createArrowFiles()* function to carry out QC and doublet removal for each sample individually(48, 121). Barcodes with at least 1,000 unique fragments, but no more than 100,000, and TSS enrichment scores greater than or equal to 8 were deemed high quality cells and were carried forward to doublet detection(45). ArchR’s *addDoubletScores()* and *filterDoublets()* functions were used to identify and remove cells predicted as doublets from further downstream analyses(48). The distributions of QC metrics, post-QC, for the patient and cell line cohort datasets are visualized in Figure S1C-D and Figure S9C-D, respectively.

### Single-cell ATAC-seq quantification, feature selection, and integration with single-cell RNA-seq

To analyze the scATAC-seq cells in the patient cohort analysis, we quantified Tn5 insertion counts in a matrix of contiguous genomic tiles 500 bp in size using ArchR’s *addTileMatrix()* function(48). As described previously, we used the iterative latent semantic indexing (LSI) procedure implemented in ArchR’s *addIterativeLSI()* function to reduce the dimensionality of the dataset (23, 44, 45, 48). We visualized the cells in a UMAP plot using ArchR’s *addUMAP()* function with the top 30 LSI components, as described previously (23, 48). UMAP plots were then plotted using the ggplot2 R package(121, 124). To integrate with matching scRNA-seq cells, we first calculated gene scores for scATAC-seq cells using ArchR’s *addGeneScoreMatrix()* function, as described previously (23, 48). Next, we used ArchR’s *addGeneIntegrationMatrix()* function to transfer cell type cluster labels and gene expression profiles from scRNA-seq cells to scATAC-seq cells, as described previously (Figure 1E; Figure S2C-E) (23, 46-48). Additionally, inferCNV status and predicted subtype labels were assigned to each scATAC-seq cell based on the annotations of its nearest neighboring cell in scRNA-seq (Figure 1G) (46-48). Using the groupList parameter in *addGeneIntegrationMatrix()*, we constrained the integration to cells from the same patient samples to ensure accurate matching of scRNA-seq and scATAC profiles(48).

The same procedures were applied to analyze the scATAC-seq cells in the Basal-like subtype, Luminal subtype, and cell line cohort analyses, with the exception of using an unconstrained integration in the cell line cohort analysis (i.e. “all versus all”) (Figure 2A-B; Figure 4A-B; Figure 6A-B). To evaluate the performance of the label transfer procedure, we leveraged the ground truth identities of the scATAC-seq cells in the cell line cohort analysis (HCC1143, SUM149PT, MCF7, or T47D) to calculate the percentage of scATAC-seq cells correctly assigned to their true cell line identity (99.71%). After transferring labels to scATAC-seq cells in the Basal-like subtype, Luminal subtype, and cell line cohort analyses, peak calling was carried out in each, as described previously, using ArchR’s *addGroupCoverages(), addReproduciblePeakSet()*, and *addPeakMatrix()* functions(13, 23, 48, 67, 68).

### Unsupervised hierarchical clustering and PCA of pseudo-bulk transcriptomes

To perform the pseudo-bulk clustering analysis, we first created pseudo-bulk transcriptomes by summing gene counts across BC cells of the majority subtype within each BC patient sample. This procedure resulted in two Basal-like-specific pseudo-bulk profiles from Patients 5 and 6, and ten Luminal-specific (Luminal A or B) pseudo-bulk profiles from Patients 7-15 (Figure S3). Similarly, we created pseudo-bulk transcriptome profiles for the normal mammary epithelial cell types from healthy controls by summing gene counts across cells of the same cell type within each healthy patient. This procedure resulted in four mature luminal, four basal epithelial, and four luminal progenitor pseudo-bulk profiles all derived from Patients 1-4 (Figure S3).

Only genes expressed across all pseudo-bulk transcriptome profiles were used for downstream analysis to avoid the possible contribution of technical zeros. The resulting matrix of 24 pseudo-bulk profiles was transformed using the regularized logarithm (rlog) transformation from the DESeq2 R package to stabilize variance and account for differences in library size between patient samples(71, 121). After this transformation, the top 10% most variably expressed genes were used for unsupervised hierarchical clustering analysis performed in the SigClust2 R package with Euclidean distance and ward.D2 as the linkage method(121, 128). This resulted in two statistically significant clusters and the dendrogram was further visualized with a heatmap of scaled pseudo-bulk profiles using the ComplexHeatmap R package (Figure S3A) (121, 129, 130). The same variably expressed genes were used for generating the PCA plots with DESeq2’s *plotPCA()* function (Figure S3B) (71).

### Differential gene expression and differential peak accessibility testing

We carried out differential gene expression testing after clustering cells within each individual patient sample, using Seurat’s *FindAllMarkers()* function with only.pos set to TRUE and test.use set to “wilcox” to identify cluster marker genes(46, 47). Genes with Bonferroni-corrected p-values <= 0.01 were deemed statistically significant marker genes.

For the comparisons of subtype-specific BC cells from BC patients to their nearest normal mammary epithelial cell types from healthy controls, we performed differential gene expression and peak accessibility testing on a pseudo-bulk scale to overcome the pseudo-replication bias in single-cell data (72, 73). Within the Basal-like subtype analysis, we created pseudo-bulk transcriptome profiles by summing gene and peak counts across Basal-like BC cells within each BC patient. The same operation was performed for normal luminal progenitor cells within each healthy patient. This resulted in two Basal-like BC-specific pseudo-bulk profiles from Patients 5 and 6 and four luminal progenitor pseudo-bulk profiles from Patients 1-4. The same procedure was used to construct pseudo-bulk transcriptome profiles within the Luminal subtype analysis, resulting in nine Luminal BC-specific pseudo-bulk profiles and four mature luminal pseudo-bulk profiles. Note that Patient 9 did not have a sufficient number of cells for the peak-to-gene association analysis (n=111) and was therefore excluded from differential gene expression and peak accessibility testing.

Within both Basal-like and Luminal subtype analyses, uninformative genes and peaks with zero counts across all pseudo-bulk profiles were removed. The resulting matrices within the Basal-like and Luminal subtype analyses were read into DESeq2 with the *DESeqDataSetFromMatrix()* function (71). To verify that potential differences in cell count between pseudo-bulk profiles would not confound differential gene expression and peak accessibility testing, we visualized PCA plots of the pseudo-bulk profiles colored by cell count using DESeq2’s *plotPCA()* function within the Basal-like and Luminal subtype analyses(71). These verification steps confirmed that differences in cell count between pseudo-bulk profiles were not associated with PCs 1 or 2 in both the Basal-like and Luminal subtype analyses.

Pseudo-bulk differential gene expression testing was performed with DESeq2’s *DESeq()* function and genes with FDR (Benjamini-Hochberg) adjusted p-values < 0.05 and absolute log2 fold changes > 0.58 were deemed statistically significant differentially expressed genes (71, 131). The same procedure and thresholds were used for pseudo-bulk differential peak accessibility testing to arrive at statistically significant differentially accessible peaks.

### Peak-to-gene association analysis

To quantify associations between peak accessibility and gene expression in the Basal-like and Luminal subtype cohorts, we first generated metacells (i.e. aggregates of 100 similar scATAC-seq cells) via a k-nearest neighbor procedure stratified by patient in the low dimensional LSI space for each cohort’s scATAC-seq analysis(48). This procedure resulted in patient-specific metacells that could be classified into the “cancer” or “normal” conditions, based on the sample of origin of constituent scATAC-seq cells within each metacell. Metacells with more than 80% overlap of cell composition with any other metacell from the same patient were removed from further downstream analysis (48). Peak accessibility and gene expression were summarized for each metacell by summing peak counts and inferred gene expression values (from the matching scRNA-seq data) across scATAC-seq cells within each metacell(48). The resulting peak and gene expression matrices were normalized to counts per 10,000 and log2-transformed with a pseudo-count of 1(48). These peak and gene expression matrices were used as input into the peak-to-gene association analyses for each patient cohort dataset. The same set of procedures was applied to the cell line cohort dataset, with the exception of stratifying the k-nearest neighbor procedure by cell line and classifying the resulting cell line-specific metacells into the “Basal-like” or “Luminal” conditions based on the sample of origin of constituent scATAC-seq cells within each metacell(48).

Within the Basal-like subtype, Luminal subtype, and cell line cohorts, we first performed independent peak-to-gene association analyses in each condition. Using the patient or cell line-specific metacells, we fit a linear mixed-effects model (LMM) with random intercepts, in each condition, using the lmerTest and lme4 R packages, to quantify the effect size of peak accessibility on gene expression for every peak located within 500 kb of each gene(53, 54, 121). More specifically, gene expression was modeled as a function of peak accessibility, treated as a fixed effect, and patient of origin, treated as a random effect to account for variation in gene expression between patients. The Satterthwaite approximation of degrees of freedom, implemented in the lmerTest R package, was used to determine statistical significance of the fixed effect term in each peak-gene model tested(53, 54, 121). To correct for multiple testing in each condition, the Benjamini-Hochberg method was applied using the *p*.*adjust()* function from the stats R package and peak-gene pairs with FDR-adjusted p-value < 1e-04 were deemed statistically significant peak-to-gene associations (Figure 2D; Figure S6A; Figure 6C) (121, 131). These were the final peak-to-gene association analyses performed for the cell line cohort dataset. However, for the Basal-like and Luminal subtype cohorts, intronic or distal intergenic peak-gene pairs with a significant association in at least one condition (“cancer” or “normal”) were carried forward to a second phase of peak-to-gene association analyses to identify changes in peak-to-gene effect size between conditions.

To quantify the change in peak-to-gene effect size between conditions in a differential peak-to-gene association analysis, we combined the patient-specific metacells from both conditions and fit another LMM with an interaction term(53, 54). More specifically, gene expression was modeled as a function of peak accessibility, condition, the interaction between peak accessibility and condition, and patient of origin. All terms were treated as fixed effects, except for patient of origin which was treated again as a random effect to account for variation in gene expression between patients. As performed in the first phase of peak-to-gene association analyses, the Satterthwaite approximation of degrees of freedom was used to determine statistical significance of the fixed effect terms in each peak-gene model tested and the resulting p-values for the interaction term were corrected for multiple testing using the Benjamini-Hochberg method implemented in the *p*.*adjust()* function from the stats R package(53, 54, 121, 131). Peak-to-gene associations with FDR-adjusted p-value < 1e-04 for the interaction term were deemed statistically significant differential peak-to-gene associations (Figure 2E-G; Figure S6B-D).

Within the Basal-like and Luminal subtype analyses, significant differential peak-to-gene associations were visualized in a scatter plot of effect sizes for each condition using ggplot2 (Figure 2E-G; Figure S6B-D) (124). The cancer-specific peak-to-gene associations involving genes upregulated in the cancer condition were visualized in a heatmap of effect sizes for each condition using the *Heatmap()* function from ComplexHeatmap (Figure 3A; Figure 4C) (129, 130). Select cancer-specific peak-to-gene associations of interest were visualized in a genomic browser track format, using ArchR’s *plotBrowserTrack()* function, to display the ATAC-seq coverage patterns of the surrounding locus stratified by condition and by patient (Figure 3C; Figure S4; Figure 4E; Figure S7) (48). The same cancer-specific peak-to-gene associations of interest were visualized in scatter plots of peak accessibility by the inferred level of gene expression in metacells using ggplot2 (Figure 3D; Figure 4F) (124).

### Overlap analyses of genomic coordinates

To identify peak-to-gene associations that overlapped with existing ENCODE annotations downloaded from https://screen.encodeproject.org, the genomic coordinates of the peaks participating in the peak-to-gene associations were converted into a *GRanges* object using the GenomicRanges R package(69, 70, 121, 132). The genomic coordinates of the existing ENCODE annotations were also converted into a second *GRanges* object(132). These two *GRanges* objects were used as input into the *subsetByOverlaps()* function from the IRanges R package(121, 132). The output from this function was a *GRanges* object containing the genomic coordinates of peaks that overlapped with the genomic coordinates of existing ENCODE annotations and was used to annotate the initial set of peak-to-gene associations for overlap with existing ENCODE annotations (Figure 2D; Figure S6A; Figure 6C).

To perform the overlap analyses of putative enhancers between *in vitro* and *in vivo* BC cells for each subtype, the genomic coordinates of the putative enhancers *in vitro* and *in vivo* were converted into *GRanges* objects(132). These two *GRanges* objects were used as input into IRanges’ *findOverlaps()* function to identify the number of overlapping, or shared, putative enhancers between *in vitro* and *in vivo* BC cells(132). The same *GRanges* objects were used as input into IRanges’ *subsetByOverlaps()* function, with the invert parameter set to TRUE, to identify the numbers of putative enhancers specific to *in vitro* and *in vivo* BC cells (Figure 6G; Figure 6N) (132).

To perform the overlap analyses of putative enhancer-target gene pairs between *in vitro* and *in vivo* BC cells for each subtype, the genomic coordinates ofthe putative enhancers participating in the putative enhancer-target gene pairs *in vitro* and *in vivo* were converted into *GRanges* objects(132). These two *GRanges* objects were used as input into IRanges’ *findOverlaps()* function to return a *GRanges* object containing the indices of genomic coordinates that overlapped between the two input *GRanges* objects(132). The indices of overlapping genomic coordinates from each *GRanges* object were used to merge both sets of putative enhancer-target gene pairs, *in vitro* and *in vivo*, into one matrix based on overlapping putative enhancers. These sets of putative enhancer-target gene pairs, *in vitro* and *in vivo*, were screened for those with overlapping putative enhancers that linked to the same gene in both *in vitro* and *in vivo* settings. The resulting set of shared putative enhancer-target gene pairs in both settings was used to annotate the overlap status for the initial sets of putative enhancer-target gene pairs identified in each setting (Figure 6H; Figure 6O).

### Gene set enrichment analysis

To perform gene set enrichment analysis with Hallmark gene sets from MSigDB, we inputted the list of genes of interest into the *enricher()* function from the clusterProfiler R package to test for significant enrichments via hypergeometric tests (74-76, 121). Gene sets with Benjamini-Hochberg adjusted p-values < 0.05 were deemed statistically significant enrichments(131). The top three most significantly enriched gene sets were visualized using the *dotplot()* function from the enrichplot R package with the showCategory parameter set to “3” (Figure 3B; Figure 4D; Figure 6F; Figure 6M) (133).

### Survival analysis

CALGB 40603 clinical data were acquired from dbGaP (phs001863.v1.p1) and upper quartile normalized RNA-seq expression data were downloaded from GEO (GSE154524). The normalized expression values were further log2-transformed.

FUSCC clinical data were downloaded from Table S1 of Jiang et al 2019 and RNA-seq fastq files were acquired from the NCBI Sequence Read Archive (SRP157974)(134). RNA-seq fastqs were aligned to the hg38 reference genome with STAR 2.7.6a and quantified with Salmon 1.4.0. Salmon counts were then upper quartile normalized and log2(x+1) transformed.

METABRIC clinical data and normalized expression data were acquired from the European Genome-Phenome Archive (EGAS00000000083) and came from Curtis et al 2012(135). The normalized expression values were further log2(x+1) transformed. PR IHC data and HER2 FISH data were missing from this dataset, so TNBC samples were defined as IHC ER negative, no HER2 SNP6 gain, and HER2 SNP6 loss or HER2 negative by expression. Correspondingly, HR+/HER2-samples were defined as IHC ER positive, no HER2 SNP6 gain, and HER2 SNP6 loss or HER2 negative by expression. Stage 0, stage 4, and untreated samples were excluded from analysis.

SCAN-B clinical data and gene-level FPKM RNA-seq expression data were downloaded from Mendeley Data (https://data.mendeley.com/datasets/yzxtxn4nmd/3) and came from Staaf et al 2022(136). The FPKM expression values were further upper quartile normalized and log2(x+1) transformed. Untreated samples, bilateral samples, multi-centric samples, lymph node samples, and normal samples were excluded from the analysis. Furthermore, sample duplicates were excluded (keeping the specimen that had the most frequently used library protocol, sequencer serial, library barcode, or pool name in the dataset, respectively).

TCGA-BRCA clinical data and RNA-seq fastqs were acquired from the Genomic Data Commons Data Portal (https://portal.gdc.cancer.gov/projects/TCGA-BRCA)(137). RNA-seq fastqs were aligned to the hg38 reference genome with STAR 2.7.6a and quantified with Salmon 1.4.0. Salmon counts were then upper quartile normalized and log2(x+1) transformed. Only fresh frozen tumor samples were considered for analysis, and stage 4 samples were excluded.

Cox proportional hazards models were fit using the survival R package for each of the 829 unique genes participating in 7,167 cancer-specific peak-to-gene associations, identified in the Basal-like subtype analysis, over the set of TNBC patients in each dataset. The same was performed for each of the 288 unique genes participating in 1,931 cancer-specific peak-to-gene associations, identified in the Luminal subtype analysis, over the set of HR+/HER2-samples in each dataset. Separate models were fit using each definition of survival available for a dataset. The survival R package was used for the Cox proportional hazard modeling with the formula Surv(time, event) ∼ gene_expression.

Kaplan-Meier plots were created using the survminer R package, using median gene expression values as the “high expression” vs. “low expression” cutoffs, with log-rank P-values displayed.

### Landscape plots

TCGA-BRCA GISTIC2.0 gene-level copy number data (all_data_by_genes.txt) were downloaded from TCGA Firebrowse (http://firebrowse.org/?cohort=BRCA#)(137). Gene-level copy number scores were then converted to 534 pre-determined chromosomal segment copy-number scores by calculating the mean score of genes overlapping a given segment. The 534 segments used included chromosomal regions associated with cancer and whole-arm chromosome segments, as described in detail in Xia et al 2019 (138). Segment-level copy number scores > 0.3 in a sample were considered copy number gains, and segment-level copy number scores < −0.3 in a sample were considered copy number losses. The percentage of all TNBC TCGA samples with a copy number gain/loss call for each segment was calculated and plotted by the relative segment order, with *HEY1* labeled between the two non-whole-arm segments it was closest to (chr8:62174237-62716885.BeroukhimS5.amp and chr8:81242335-81979194.BeroukhimS2.8q21.13.amp). Correspondingly, the percentage of all HR+/HER2-TCGA samples with a copy number gain/loss call for each segment was calculated and plotted by the relative segment order, with *CRABP2* labeled between the two non-whole-arm segments it was closest to (chr1:151026302-152973244.BeroukhimS5.amp and chr1:158317017-159953843).

### CRISPR dependency and expression analyses of BC cell lines

CRISPR dependency scores were sourced from the DepMap portal using the depmap and ExperimentHub R packages(100). CCLE RNA-seq expression data was also sourced from the depmap and ExperimentHub R packages(101). We screened for BC cell lines present in both data types before plotting dot plots of CRISPR dependency scores as well as box plots of TPM expression for *HEY1* and *CRABP2* using ggplot2 (Figure 5; Figure S8)(124).

## DATA AND CODE AVAILABILITY

□ Processed single-cell RNA-seq data and single-cell ATAC-seq will be made publicly available after publication. Raw data (10x FASTQs) will be available with controlled access via dbGaP under the accession number dbGaP: phs003253.v1.p1 (https://www.ncbi.nlm.nih.gov/gap/).
□ All code used for the presented analyses is publicly available at the GitHub repository https://github.com/RegnerM2015/scBreast_scRNA_scATAC_2024.
□ Any additional information required to reanalyze the data reported in this paper is available from the lead contact.

## ACKNOWLEDGEMENTS

We thank the patients and their families for their generous donations. We thank the University of North Carolina (UNC) Tissue Procurement Facility and UNC Translational Genomics Core Facility for helping us with procuring patient specimens and sequencing genomic libraries. We thank Michele Hayward, Stephanie Metzen, and Matt Soloway at the Office of Genomics Research for help in navigating the institutional review board (IRB) protocols and data submission process to dbGaP and GEO. Finally, we thank members of the Franco and Perou labs, for their helpful comments, suggestions, and discussions. This work was supported by grants from the NIH/National Cancer Institute (R01CA273444-03), the Susan G. Komen Breast Cancer Research Foundation (CCR19608601) the Department of Defense CDMRP Breast Cancer Research Program (BC180450) to H.L.F. Additional support is provided by the UNC Breast Cancer SPORE program (5-P50-CA058223-25) to H.L.F and C.M.P. Dr. Philip Spanheimer is supported by the National Institutes of Health grant K08CA280388.

## AUTHOR CONTRIBUTIONS

H.L.F. and C.M.P. conceived and supervised the study. Patient enrollment and specimen procurement was led by P.M.S. K.W., S.G.-R., and A.T. carried out tissue specimen collection, developed the tissue dissociation protocol, and generated scRNA-seq and scATAC-seq libraries. M.J.R. designed and performed the computational analyses with input from A.T., S.G.- R., R.M.G, J.S.P., C.M.P, and H.L.F. The manuscript was written by M.J.R. and H.L.F. with input from all authors.

## SUPPLEMENTAL FIGURES

**Figure S1.**
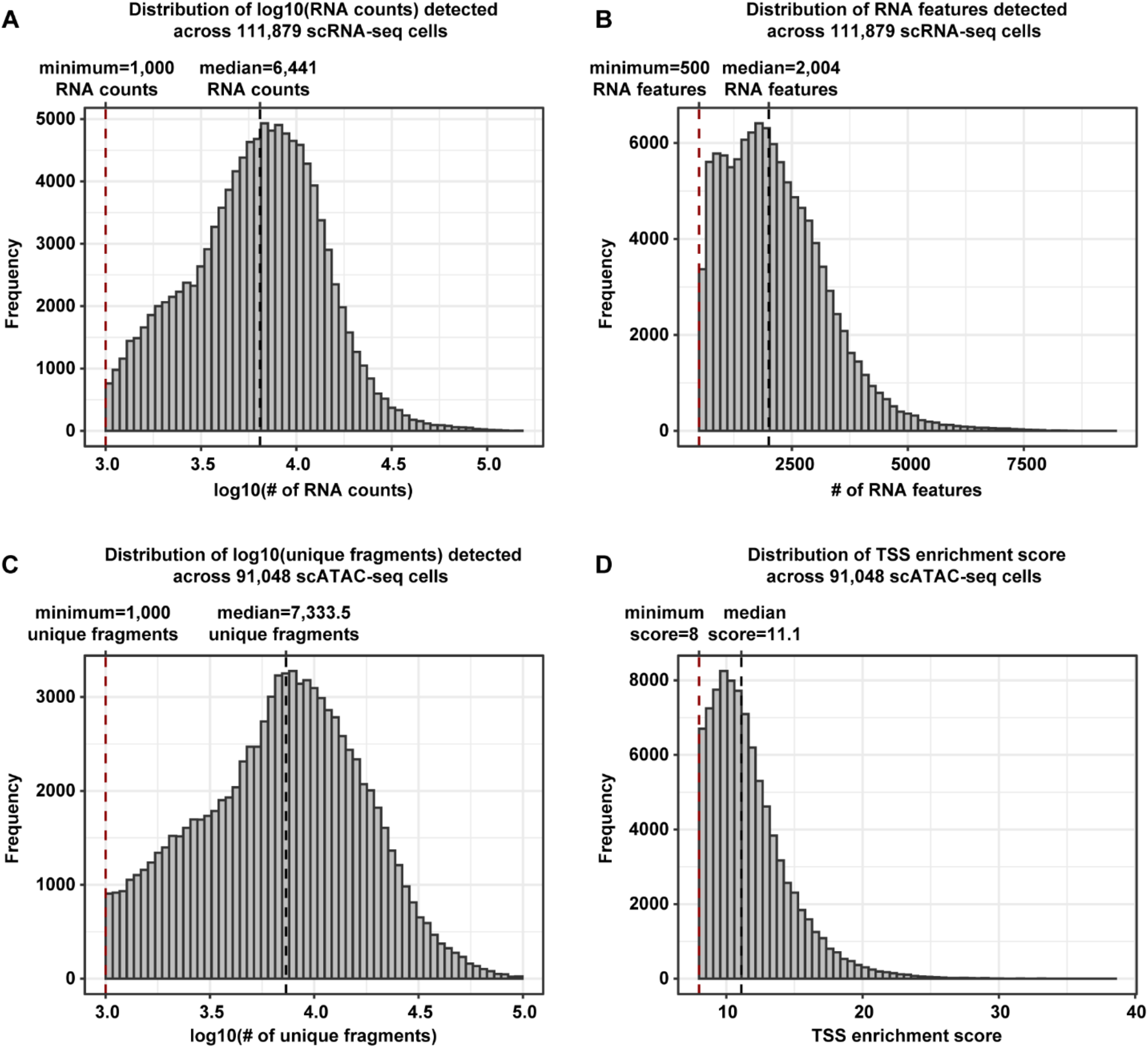
Distributions of quality control metrics for scRNA-seq and scATAC-seq cells from patient samples. **A)**. Histogram of log10(RNA counts) detected across 111,879 scRNA-seq cells from 16 patient samples. The dashed red and black lines denote the minimum (1,000 RNA counts) and median (6,441 RNA counts) number of RNA counts, respectively. **B)**. Histogram of RNA features detected across 111,879 scRNA-seq cells from 16 patient samples. The dashed red and black lines denote the minimum (500 RNA features) and median (2,004 RNA features) number of RNA features, respectively. **C)**. Histogram of log10(unique fragments) detected across 91,048 scATAC-seq cells from 16 patient samples. The dashed red and black lines denote the minimum (1,000 unique fragments) and median (7,333.5 unique fragments) number of unique fragments, respectively. **D)**. Histogram of TSS enrichment score across 91,048 scATAC-seq cells from 16 patient samples. The dashed red and black lines denote the minimum (8) and median (11.1) TSS enrichment scores, respectively.

**Figure S2.**
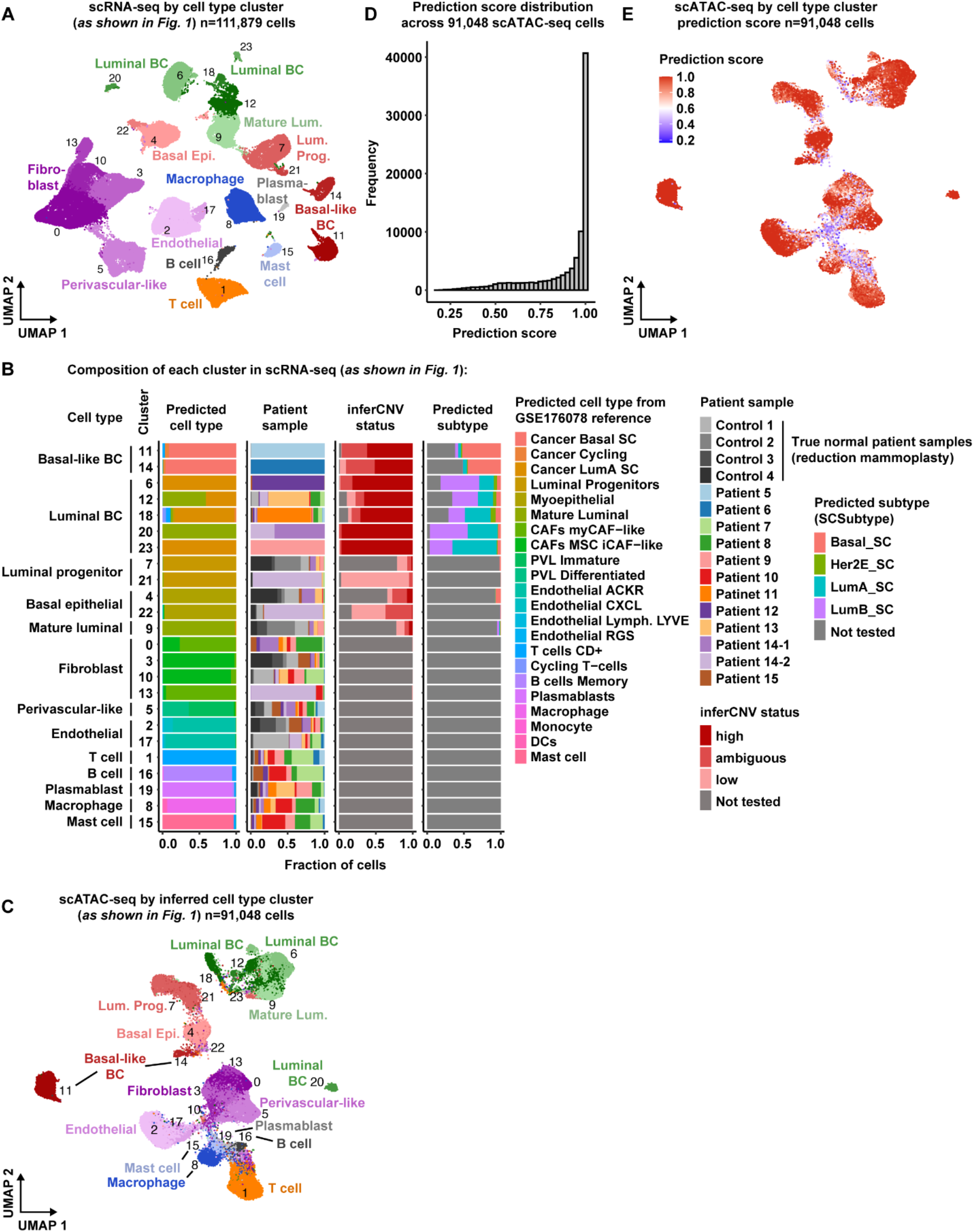
Expanded view of matched scRNA-seq and scATAC-seq for BC and healthy patient samples. **A)**. UMAP plot of 111,879 scRNA-seq cells color-coded by cell type and labeled by cluster number across 16 patient samples. **B)**. Proportion bar charts showing the composition of each cluster in scRNA-seq in terms of predicted cell type (*left*), patient sample (*middle, left*), inferCNV status (*middle, right*), and predicted subtype (*right*). Color-coded legends are shown on the right. Note that the reference scRNA-seq dataset from GSE176078 was used to help annotate clusters in each individual patient analysis. Therefore, the predicted cell type labels shown here were derived from the cluster labels annotated in each individual patient dataset (Methods). **C)**. UMAP plot of 91,048 scATAC-seq cells color-coded by inferred cell type and labeled by inferred cluster number across 16 patient samples. **D)**. Histogram of cell type cluster prediction scores across 91,048 scATAC-seq cells. **E)**. UMAP plot of scATAC-seq cells, as shown in panel **C**, but colored by cell type cluster prediction score.

**Figure S3.**
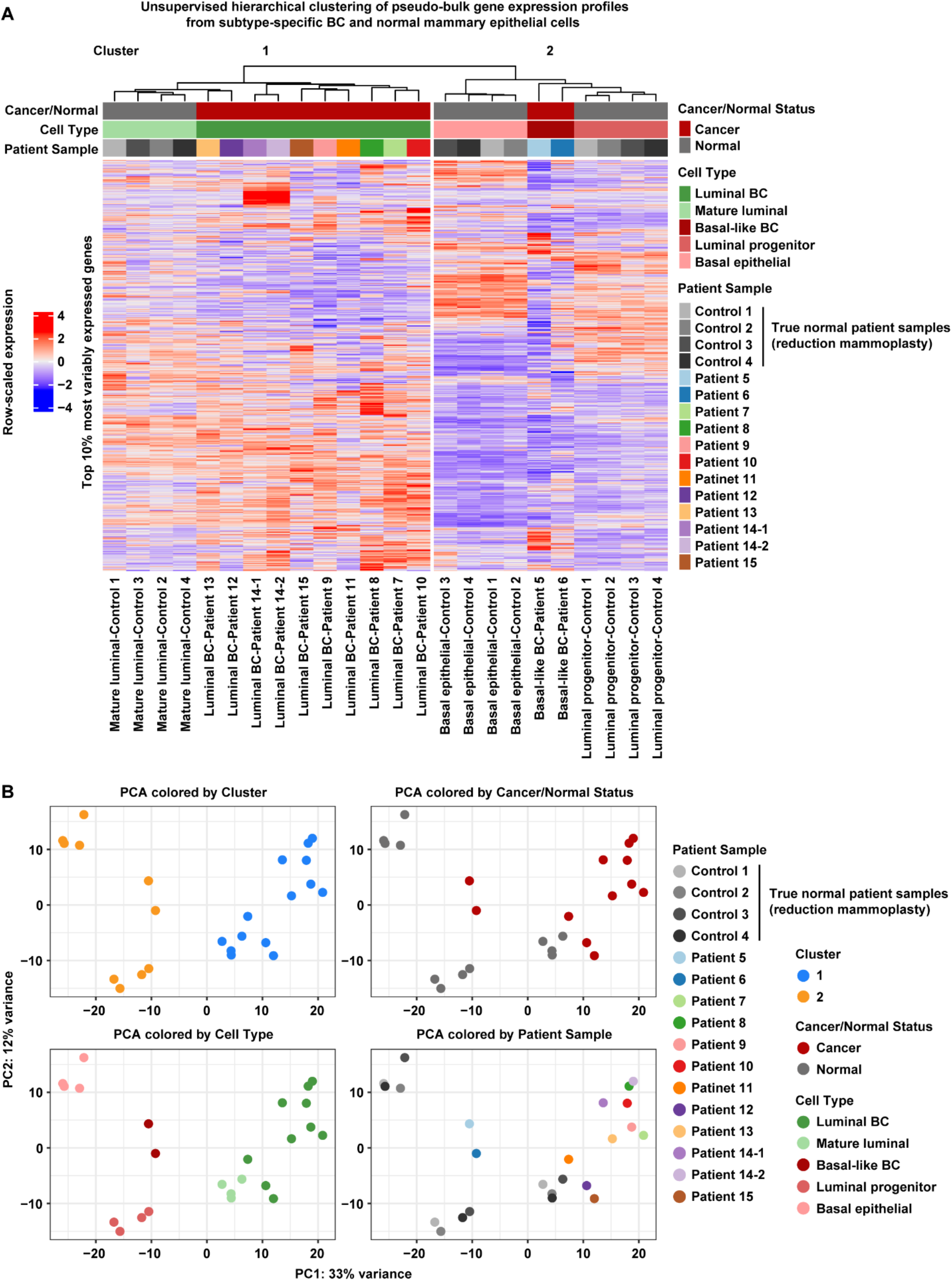
Cluster analysis of pseudo-bulk transcriptomes from subtype-specific BC cells and normal mammary epithelial cells from healthy controls. **A)**. Unsupervised hierarchical clustering and row-scaled heatmap of pseudo-bulk transcriptomes (*columns*) using the top 10% most variably expressed genes (822 genes) (*rows*). The indicated clusters in the dendrogram are statistically significant as determined by SigClust2 (Methods). The specific groups of single cells used to create each pseudo-bulk transcriptome are annotated above and below the heatmap. Color-coded legends for each set of annotations are shown to the right. **B)**. Principal Component Analysis (PCA) of pseudo-bulk transcriptomes, as shown in panel **A**, using the top 10% most variably expressed genes (822 genes). Pseudo-bulk transcriptomes are color-coded by SigClust2 cluster as shown in panel **A** (*top left*), cancer/normal status (*top right*), cell type (*bottom left*), and patient sample (*bottom right*). Color-coded legends for each set of annotations are shown to the right.

**Figure S4.**
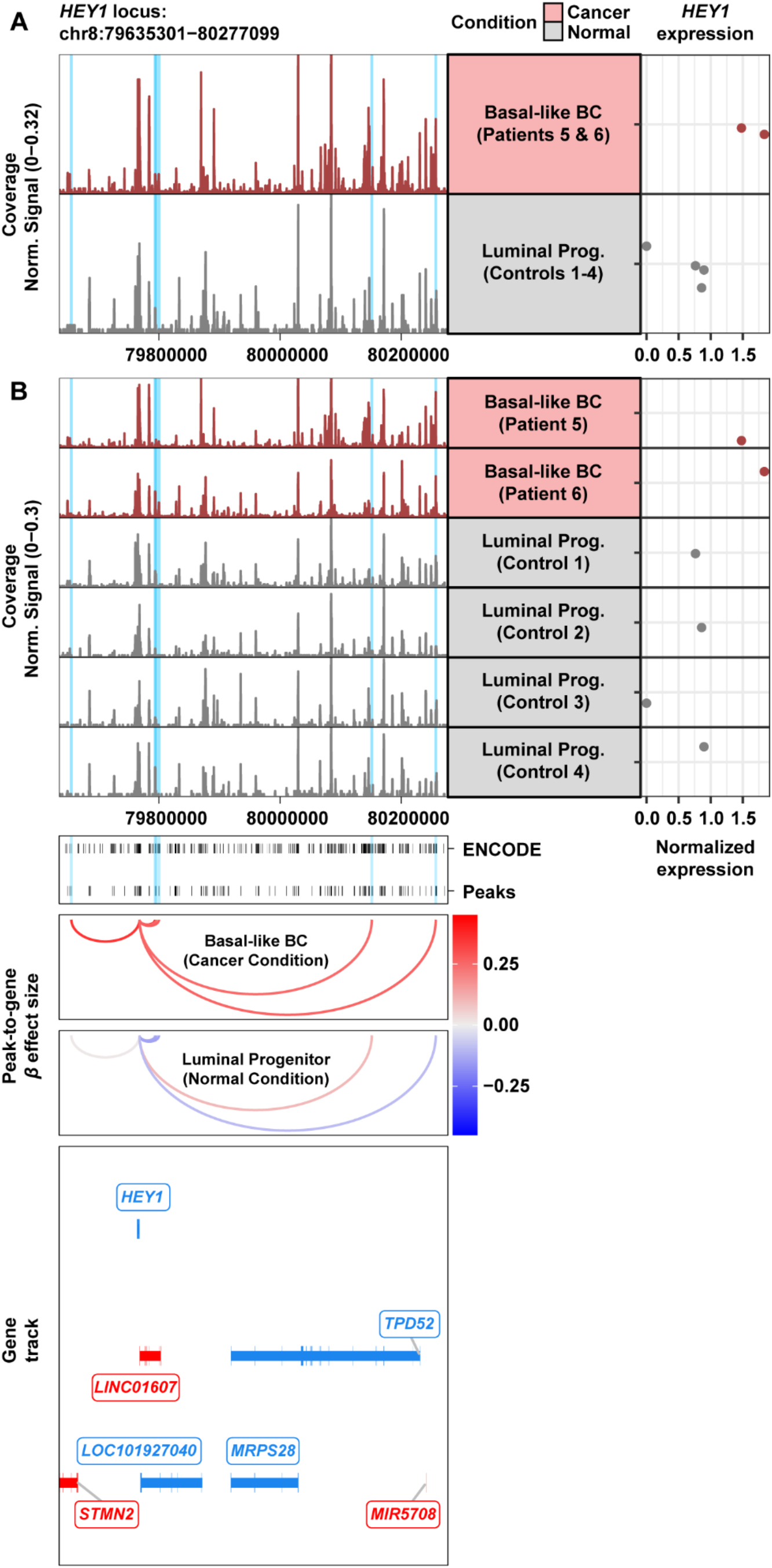
Expanded view of putative cancer-specific enhancers linked to *HEY1* expression. **A)**. Browser track showing the accessibility profile at the *HEY1* locus stratified by condition (*top*). The putative cancer-specific enhancers linked to *HEY1* expression are highlighted in light blue. Matching pseudo-bulk scRNA-seq expression of *HEY1* is shown for each condition (*right*). **B)**. Browser track, as in panel **A**, but stratified by patient (*top*). The putative cancer-specific enhancers linked to *HEY1* expression are highlighted in light blue. Matching pseudo-bulk scRNA-seq expression of *HEY1* is shown for each patient (*right*). ENCODE regulatory element annotations and peaks called from the scATAC-seq data, are shown below the browser track (*middle*). Peak-to-gene loops show the standardized effect sizes, in each condition, of chromatin accessibility at the putative cancer-specific enhancers on *HEY1* expression (*bottom*).

**Figure S5.**
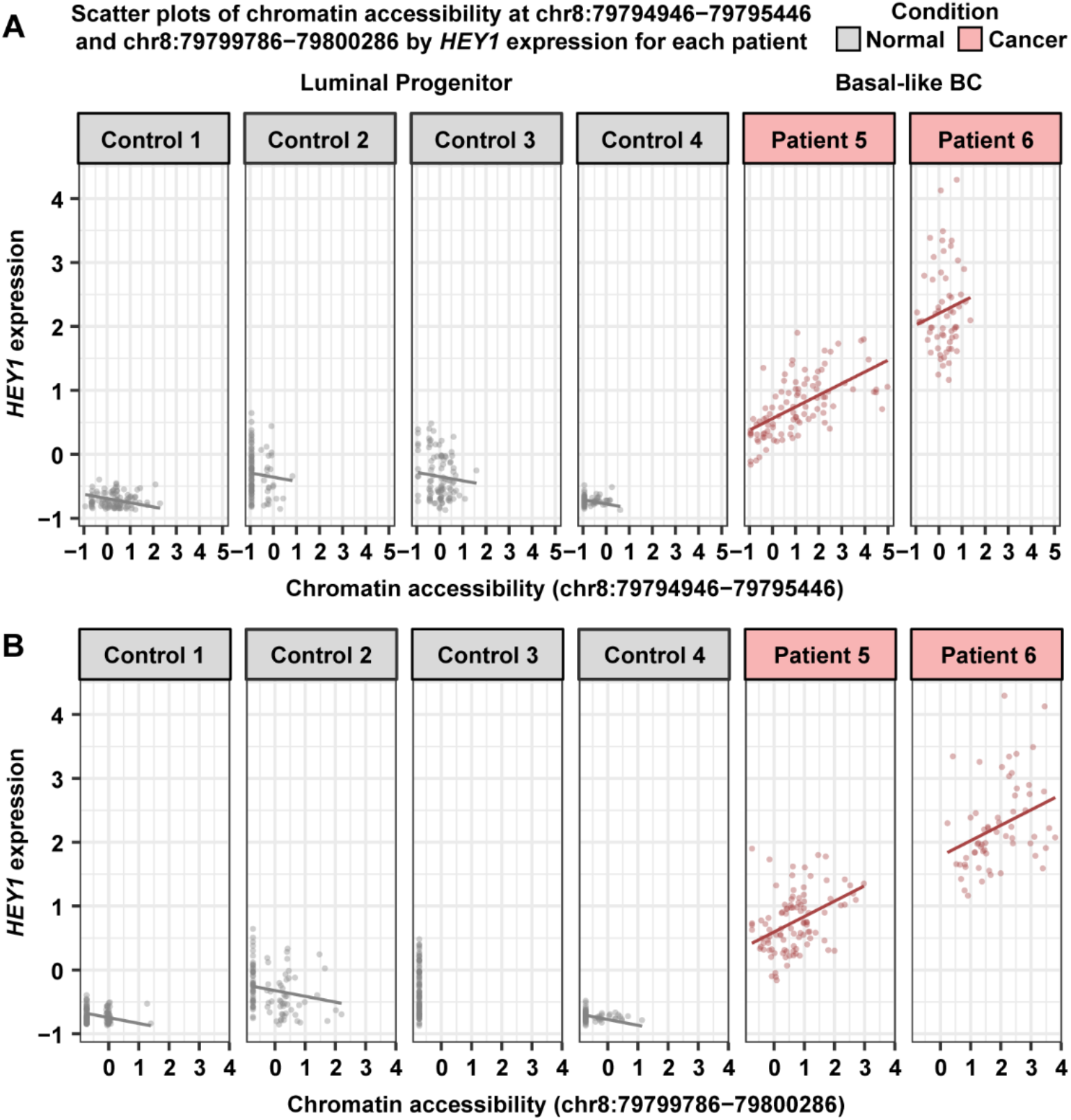
Scatter plots of chromatin accessibility by *HEY1* expression for nearest neighboring cancer-specific enhancers identified in the Basal-like subtype analysis. **A)**. Scatter plots of chromatin accessibility at the first neighboring cancer-specific enhancer by the inferred level of *HEY1* expression in scATAC-seq metacells, stratified by patient in the normal condition (*gray*) and in the cancer condition (*red*). **B)**. Scatter plots, as in panel **A**, but for the second neighboring cancer-specific enhancer.

**Figure S6.**
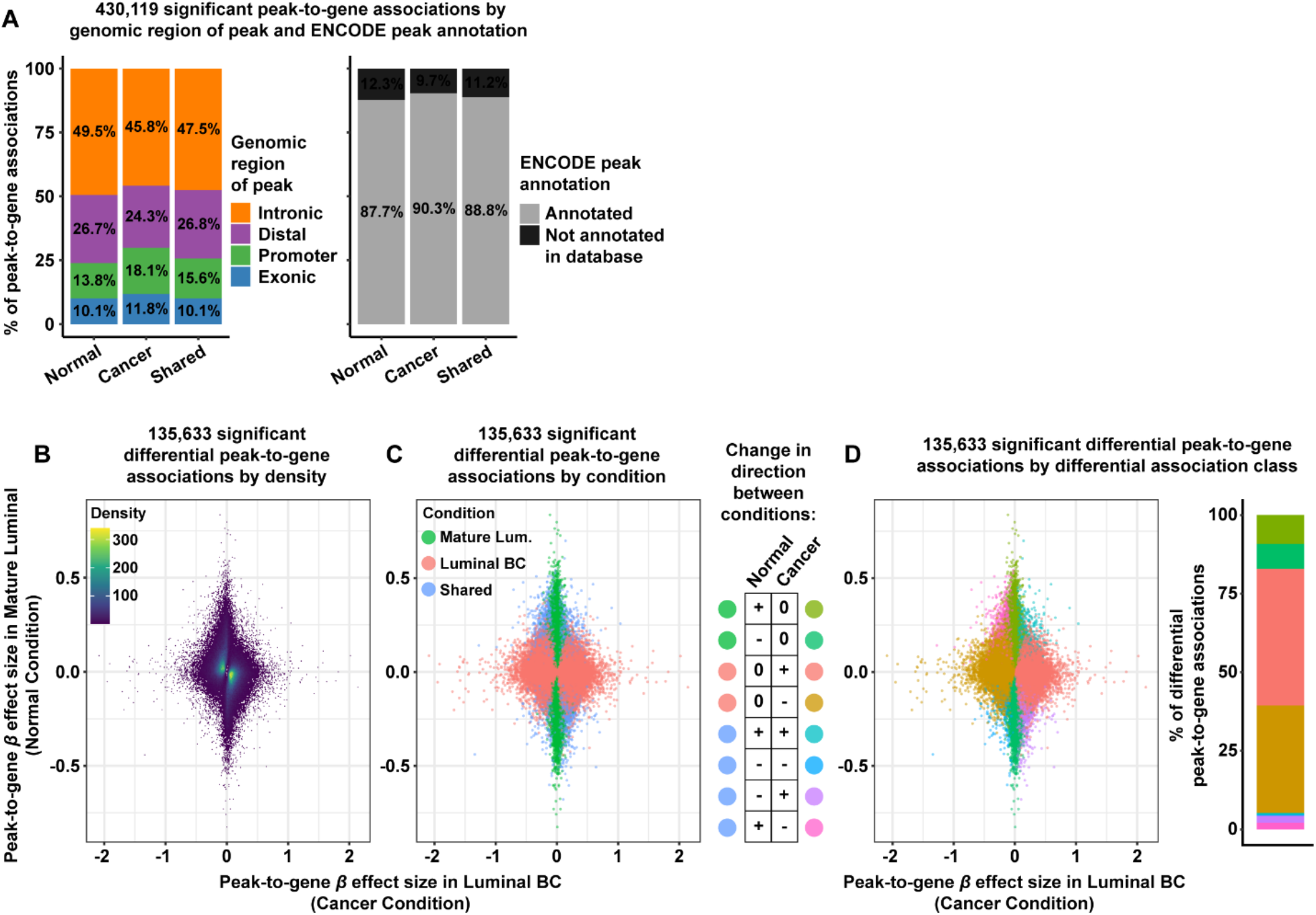
Expanded view of differential peak-to-gene associations identified in Luminal BC cells relative to mature luminal cells. **A)**. Proportion bar charts showing the genomic distribution (*left*) and ENCODE annotation status (*right*) for 64,808 normal-specific peak-to-gene associations, 347,033 cancer-specific associations, and 18,278 shared associations. **B)**. Density scatter plot showing the effect sizes of significant differential peak-to-gene associations in the cancer condition, comprised of Luminal BC cells, and in the normal condition, comprised of mature luminal cells. Each dot represents a peak-gene pair with a significant change in effect size between conditions. **C)**. Same scatter plot as in panel **B**, but colored by condition specificity (*left*). Infographic describing the possible directional changes in effect size between conditions (*right*). **D)**. Same scatter plot as in panel **C**, but colored by changes in direction of effect size between conditions (*left*). Proportion bar chart showing the distribution of differential association classes (*right*).

**Figure S7.**
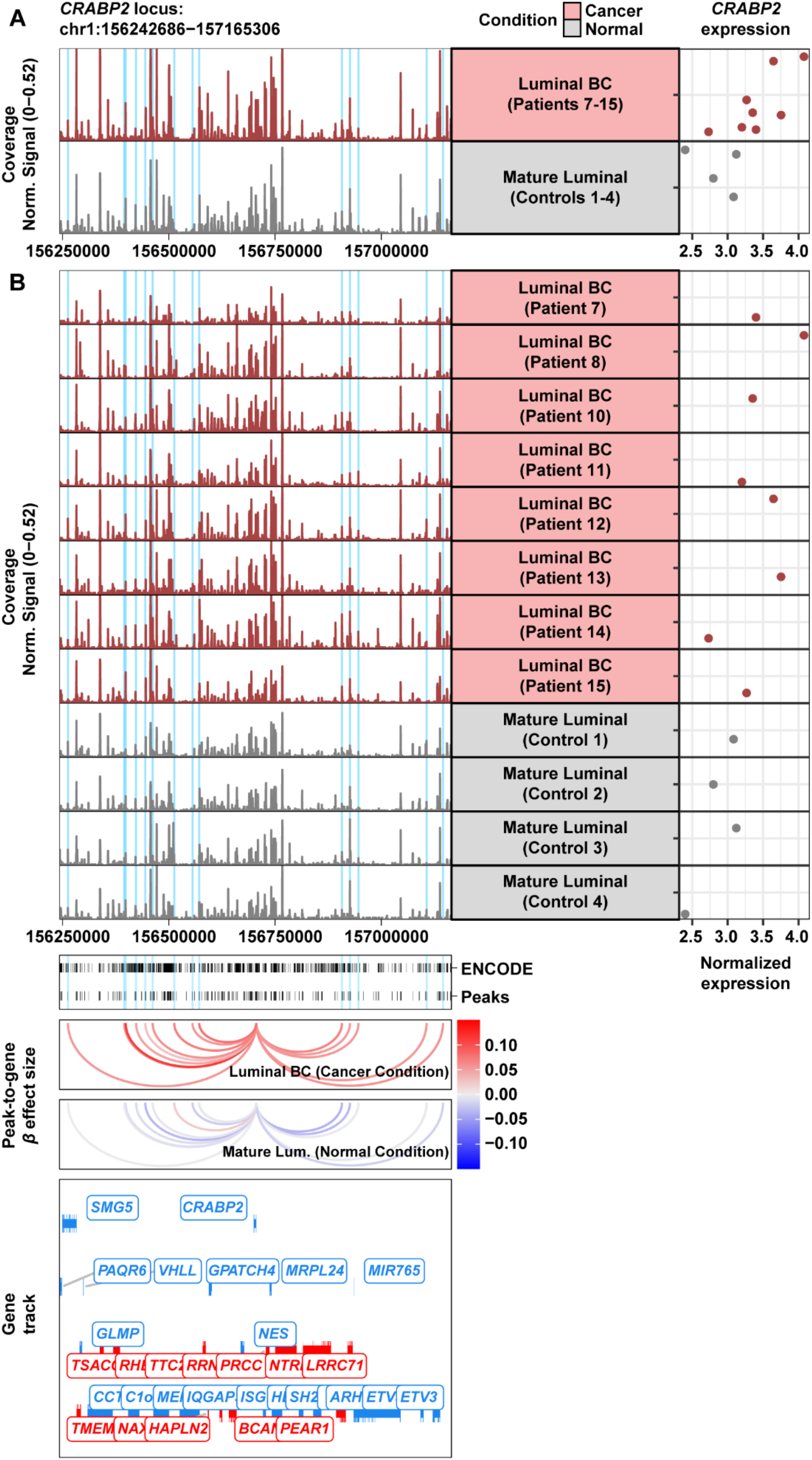
Expanded view of putative cancer-specific enhancers linked to *CRABP2* expression. **A)**. Browser track showing the accessibility profile at the *CRABP2* locus stratified by condition (*top*). Note that Patient 9 was excluded from the accessibility profile of the cancer condition, as this patient did not have a sufficient number of cells for the peak-to-gene association analysis (n=111). The putative cancer-specific enhancers linked to *CRABP2* expression are highlighted in light blue. Matching pseudo-bulk scRNA-seq expression of *CRABP2* is shown for each condition (*right*). **B)**. Browser track, as in panel **A**, but stratified by patient (*top*). The putative cancer-specific enhancers linked to *CRABP2* expression are highlighted in light blue. Matching pseudo-bulk scRNA-seq expression of *CRABP2* is shown for each patient (*right*). ENCODE regulatory element annotations and peaks called from the scATAC-seq data, are shown below the browser track (*middle*). Peak-to-gene loops show the standardized effect sizes, in each condition, of chromatin accessibility at the putative cancer-specific enhancers on *CRABP2* expression (*bottom*).

**Figure S8.**
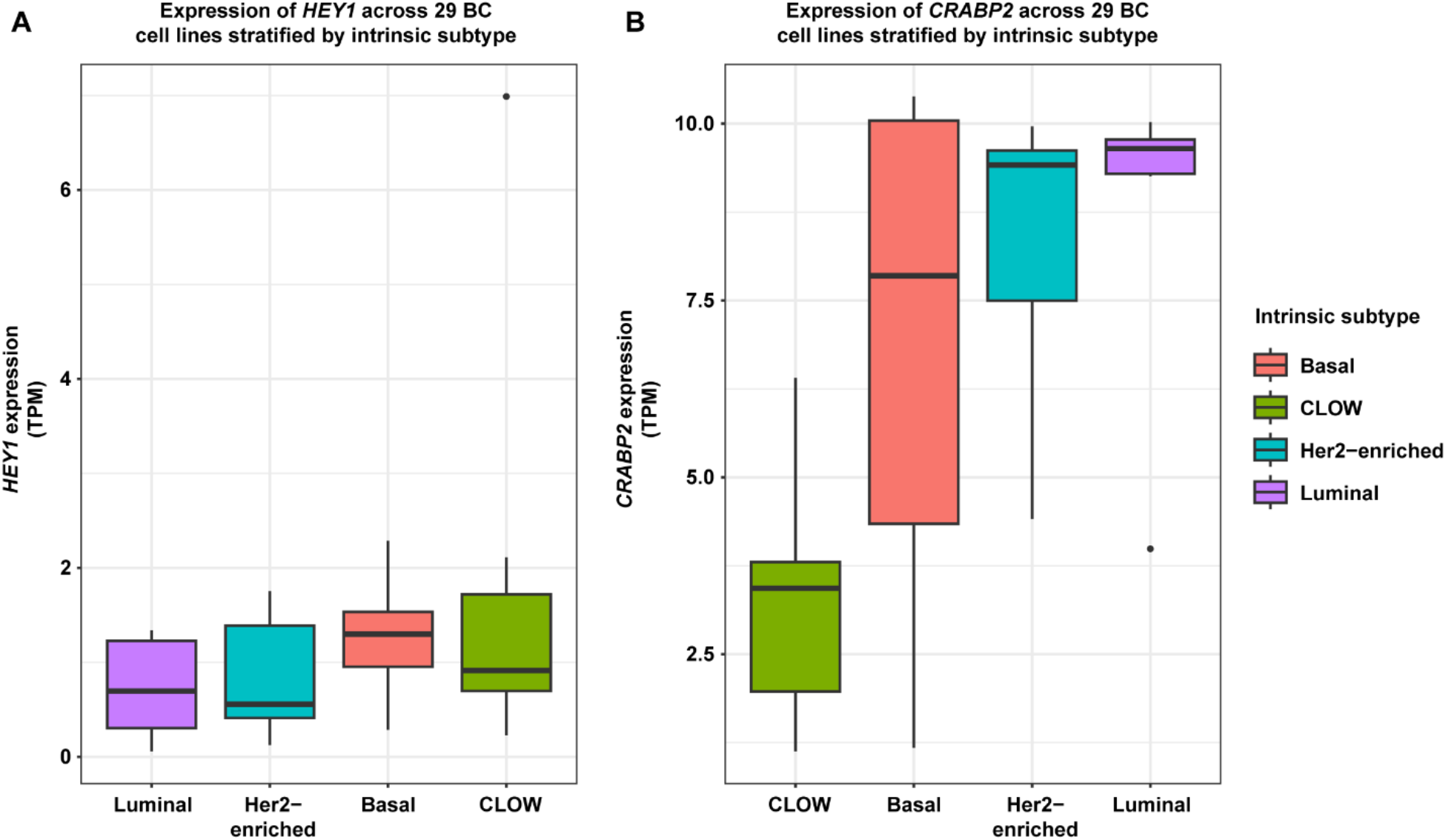
Expression of *HEY1* and *CRABP2* in 29 breast cancer CCLE cell lines stratified by intrinsic subtype. **A)**. Box plots showing the distribution of *HEY1* RNA-seq expression in transcripts per million (TPM) across CCLE BC cell lines stratified by intrinsic subtype. **B)**. Box plots, as in panel **A**, but for *CRABP2* expression.

**Figure S9.**
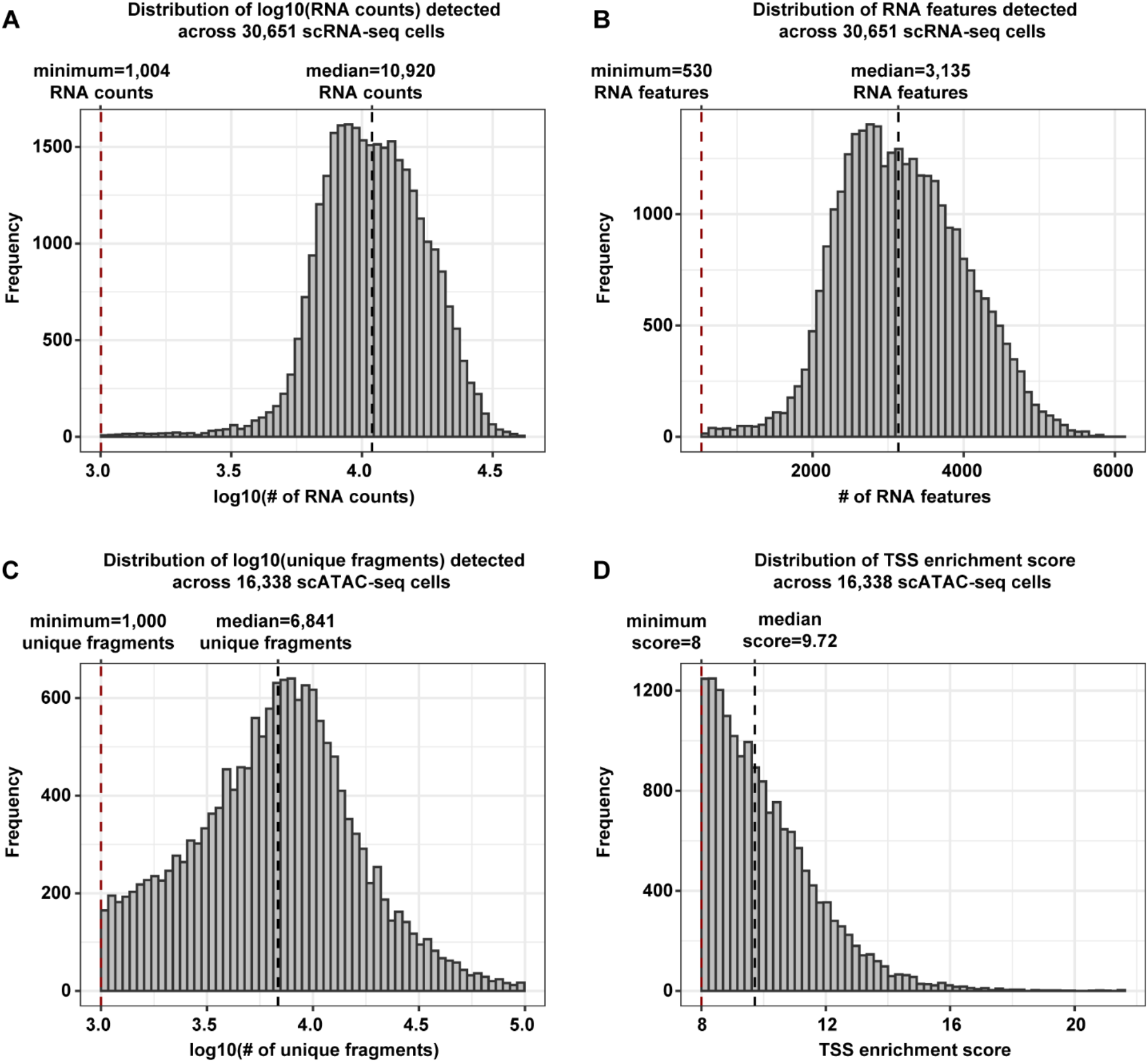
Distributions of quality control metrics for scRNA-seq and scATAC-seq cells from cell line samples. **A)**. Histogram of log10(RNA counts) detected across 30,651 scRNA-seq cells from 4 cell line samples. The dashed red and black lines denote the minimum (1,004 RNA counts) and median (10,920 RNA counts) number of RNA counts, respectively. **B)**. Histogram of RNA features detected across 30,651 scRNA-seq cells from 4 cell line samples. The dashed red and black lines denote the minimum (530 RNA features) and median (3,135 RNA features) number of RNA features, respectively. **C)**. Histogram of log10(unique fragments) detected across 16,338 scATAC-seq cells from 4 cell line samples. The dashed red and black lines denote the minimum (1,000 unique fragments) and median (6,841 unique fragments) number of unique fragments, respectively. **D)**. Histogram of TSS enrichment score across 16,338 scATAC-seq cells from 4 cell line samples. The dashed red and black lines denote the minimum (8) and median (9.72) TSS enrichment scores, respectively.

